# Cytochrome *b*_5_ reductase 4 efficiently reduces Neuroglobin and Cytoglobin

**DOI:** 10.1101/2025.07.16.665236

**Authors:** Anthony W. DeMartino, Onaje Cunningham, Saumika Mulluri, Deandra Cassel, Megan Hutar, Lawrence Ji, Efthimios A. Deligiannidis, Stefan J. Kaliszuk, Xueyin N. Huang, Qin Tong, Yvette Y. Yien, Matthew R. Dent, Jason J. Rose, Jesús Tejero

**Author notes:** Present address: Department of Medicine, University of Maryland School of Medicine, Baltimore, MD 21201, USA. Present address: Department of Chemistry, Wayne State University, Detroit, MI 48202. USA. To whom correspondence should be addressed: Jesús Tejero, Heart, Lung, Blood and Vascular Medicine Institute, University of Pittsburgh. E1246 Biomedical Science Tower, 200 Lothrop Street, Pittsburgh PA 15261.

## Abstract

Cytoglobin and Neuroglobin are heme-containing proteins expressed in most vertebrates, including mammals, with still not completely defined physiological roles. Most of the putative functions of cytoglobin/neuroglobin, such as oxygen binding or nitric oxide dioxygenation, rely on the heme iron being in the ferrous (Fe^2+^) oxidation state. Therefore, it is very possible that reducing systems are active in the cell to maintain both proteins in the ferrous state. We have previously shown that the cytochrome *b*_5_ reductase isoform 3/ cytochrome *b*_5_ system, the canonical reductase of hemoglobin and myoglobin, can reduce cytoglobin at very fast rates, consistent with a possible physiological role. However this reducing system is unable to reduce neuroglobin, which to date lacks a validated, physiologically feasible reducing system.

Here we have studied the interaction of cytochrome *b*_5_ reductase isoform 4 with cytoglobin and neuroglobin and found that cytochrome *b*_5_ reductase 4 can reduce cytoglobin at rates comparable to those observed with cytochrome *b*_5_ reductase 3/ cytochrome *b*_5_. Remarkably, it can also reduce neuroglobin efficiently. Studying different surface mutations of cytoglobin and neuroglobin we note that some cytoglobin mutations, in particular R84E and K116E decrease reduction rates by more than 10-fold, whereas surface mutations in neuroglobin that were shown to impair the interaction of neuroglobin with cytochrome *c* (E60K/D73K/E87K) show little effect on the reduction rates. We conclude that cytochrome *b*_5_ reductase 4 can supplement cytochrome *b*_5_ reductase 3/ cytochrome *b*_5_ roles for cytoglobin reduction in vivo and is a strong candidate for a physiological role as neuroglobin reductase.

## INTRODUCTION

Neuroglobin (Ngb) and Cytoglobin (Cygb) are two mammalian proteins of the globin family of yet unresolved physiological functions [1–3]. Both proteins contain a heme cofactor non-covalently bound. Unlike the evolutionarily related proteins hemoglobin (Hb) and myoglobin (Mb), the heme iron in Ngb and Cygb is not penta-coordinated but hexa-coordinated, with two histidine side chains bound to the iron in both ferric and ferrous states. In Mb/Hb the distal histidine is located further from the heme and does not directly coordinate the heme, whereas ligand binding in Ngb/Cygb is dependent on the dissociation equilibrium of the distal histidine side chain. This feature translates into specific differences in properties of Ngb and Cygb as compared to other globins, noticeable in their reaction with substrates and the coordination of gaseous ligands [1–3].

The low concentration of Ngb and Cygb in cells, their high oxygen affinity and their fast autoxidation rates suggest that the concentration of the oxy-species in vivo may be too low to fulfill oxygen transport and storage functions [1–5]. A number of possible roles have been proposed for both proteins, including, but not limited to, ROS/RNS detoxification [4–6], nitrite reduction to nitric oxide (NO) [7–11] superoxide dismutase [12], protection against DNA damage [13], regulation of cilia function [14], and lipid peroxidation [15, 16]. Both proteins form intramolecular disulfide bonds (Cys46-Cys55 in Ngb; Cys38-Cys83 in Cygb) that can regulate the activity of the heme center through their disulfide/free thiol redox equilibrium [6, 7, 9, 13, 15–18]

Most reactions that may be physiologically relevant for Cygb/Ngb require a ferrous heme species, either as deoxy (Fe^2+^) or ferrous dioxygen (Fe^2+^-O_2_), which is oxidized to ferric (Fe^3+^) heme upon reaction, or by autoxidation. Therefore, the maintenance of a catalytic cycle –or the regeneration of a reduced heme species capable of oxygen delivery-would require the presence of a reducing system to regenerate the ferrous species. The identity of this system in vivo remains unclear. For Cygb, ascorbate and the cytochrome *b*_5_ reductase isoform 3 (CYB5R3)/ cytochrome *b*_5_ (CYB5) couple have been proposed as reducing systems [19–21]. We have shown that the CYB5R3/CYB5 couple provides efficient reduction of Cygb in vitro [22], and this ability is conserved through vertebrate evolution, as shown by the reactions of human and zebrafish Cygb and CYB5R3 proteins [23]. Studies in vivo have confirmed the role of CYB5R3 in Cygb reduction [19], allowing for the control of NO diffusion via the NO dioxygenase reaction of Cygb [20]. Nevertheless, it is unclear if other reductants for Cygb, apart from CYB5R3/CYB5, may be relevant in vivo, perhaps in a cell-type dependent fashion. In our previous studies, we have also shown that the CYB5R3/CYB5 system was ineffective to reduce Ngb as compared to the effective reduction of Cygb, and its bona fide substrates Mb and Hb [22]. There are indications that mouse brain and liver homogenates can reduce Ngb [24]. Some recent studies have explored the reduction of Ngb by bovine brain extracts, finding a marginal reducing activity of some fractions over a 2 hour-period; no specific reductant proteins were identified [25]. It has been shown that *E. coli* can reduce both Cygb and Ngb through the reaction with one or more NAD(P)H dependent reductases, including NADH:flavorubredoxin oxidoreductase [26]. Due to the sequence similarity between NADH:flavorubredoxin oxidoreductase and the mammalian flavoprotein AIF (Apoptosis-Inducing Factor), and the postulated roles of Ngb on the regulation of apoptosis [27], AIF was suggested as a possible reductant for Ngb. However, initial reports indicated no effect of AIF on Ngb heme reduction [28]. Thus, no physiological reductant has been yet identified for Ngb.

Cytochrome *b*_5_ reductases (CYB5Rs) are FAD-containing proteins that catalyze electron transfer from the two-electron donors NADH or NADPH to single electron acceptors via its specific redox partner, the heme protein CYB5 [29] (Figure 1). CYB5Rs contain FAD and NAD(P)H domains structurally and sequence-related to the ferredoxin-NADP^+^ reductase protein family [30]. There are 5 genes encoding CYB5Rs in humans; CYB5R1, 2 and 3 are highly similar in sequence, probably of monophyletic origin, showing slight differences in their N-terminal sequences that translate in specific cellular localization patterns [29]. CYB5R4 and CYB5RL encode two proteins with enough sequence homology to indicate their provenance from a common ancestor to the CYB5R1/2/3 clade, but also significant identity difference among themselves and with the CYB5R1/2/3 clade, to suggest different, specific roles form CYB5R1/2/3, and their functions are largely unknown [29](Supplemental Figure 1).

**Figure 1.**
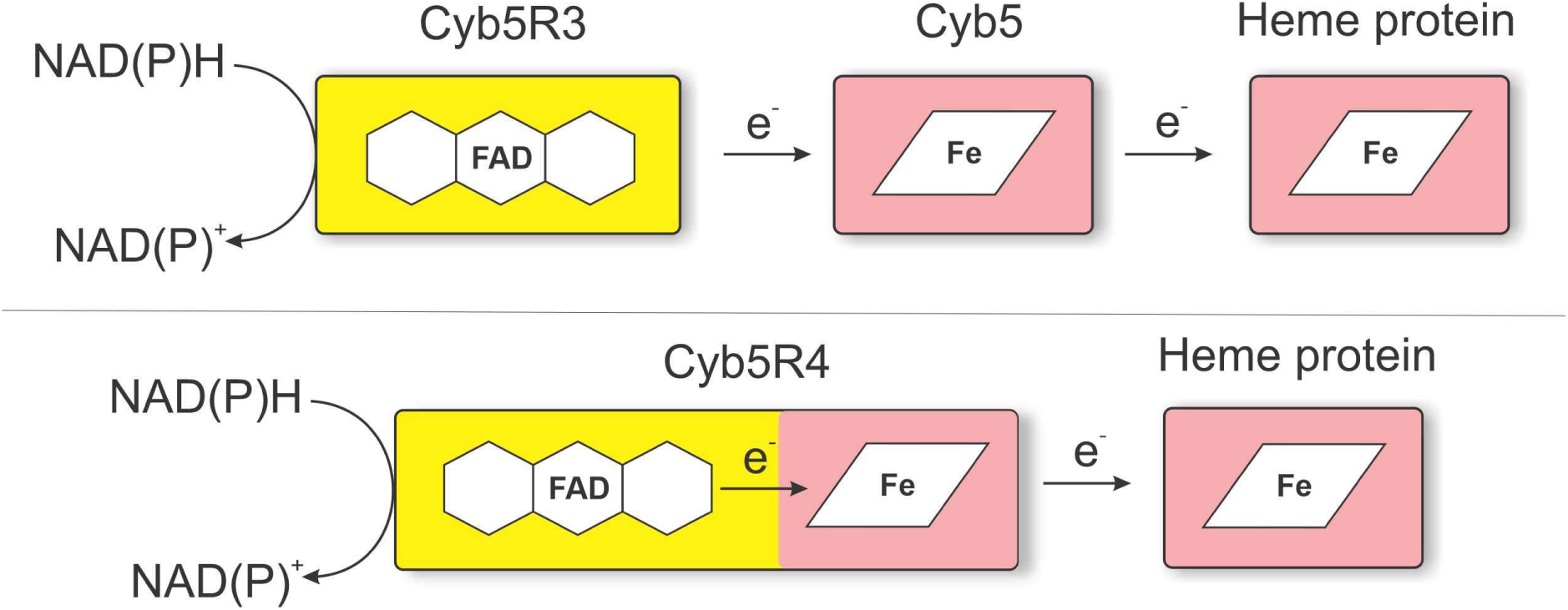
Domain arrangement and electron flow in CYB5R systems. CYB5R systems couple the two-electron donors NADH or NADPH with one-electron acceptor heme proteins. Top, the standard CYB5R hemoprotein reducing system comprises two distinct proteins; a flavoprotein reductase (yellow), which accepts electrons from NADH/NADPH to reduce its FAD cofactor; and a one-electron carrier heme protein (Cyb5, red) which receives one electron at a time from the reductase FAD. This cytochrome *b*_5_ heme protein transfers electrons to other heme proteins, such as cytoglobin. CYB5R4 (bottom) is a fusion protein including both flavoprotein reductase and heme domain fused together in a single polypeptide. Note that despite the arrangement shown in the figure, the *b*_5_ domain of CYB5R4 is located at the N-terminus of the sequence.

In this work we have focused on the reaction of CYB5R4 with Ngb and Cygb. CYB5R4, unlike the other 4 CYB5R isoforms, does not require a separate CYB5 protein as intermediate acceptor. Instead, the CYB5R4 protein carries a fused CYB5 domain in its N-terminus [31](Figure 1). In addition, this CYB5 domain has a more negative redox potential (−108 mV,[32]) than that of the common CYB5R partners CYB5a and CYB5b (−28mV to +13 mV for CYB5a, –102 to –40mV for CYB5b) [23, 32]. Our results indicate that CYB5R4 can reduce effectively Cygb and Ngb, with reduction rates higher than those observed for putative substrates such as Mb or Hb. Moreover, this activity allows for the maintenance of catalytic NO dioxygenase activity for both Cygb and Ngb in vitro. Altogether, we show that CYB5R4 can function as a Ngb/Cygb reductase complementing the reducing function of CYB5R3/CYB5 for Cygb and constitutes the first functional reducing system identified for Ngb.

## MATERIALS AND METHODS

### Reagents

All reagents were purchased from Sigma-Aldrich (St. Louis, MO) unless otherwise specified.

### Expression and Purification of Recombinant Proteins

Human Cygb was expressed recombinantly in *Escherichia coli* as previously described [10, 16, 22]. Mutation L46H in the human Cygb gene were introduced using the QuikChange site-directed mutagenesis kit (Stratagene, La Jolla, CA) using adequate primers. The Cygb containing an N-terminal Histag was purified by Ni-nitrilotriacetic acid (Ni-NTA) agarose chromatography using an Akta Purifier 10 FPLC system (GE Healthcare) with UNICORN software.

Wild-type (WT) and mutant human Ngbs were expressed and purified as described [8, 33]. SoluBL21 *Escherichia coli* cells (Genlantis, San Diego, CA) carrying the pET28-Ngb plasmid were grown, harvested and lysed as described [8]. The proteins were purified by DEAE anion exchange chromatography followed by centrifugation through an Amicon Ultra centrifugal filter (Millipore) with a 50 kDa cutoff to remove high-molecular weight contaminants. The flow-through was concentrated using a 10 kDa cutoff Amicon Ultra centrifugal filter (Millipore) and buffer exchanged to 100 mM phosphate buffer (pH 7.4) as previously described [8].

WT human CYB5R3 (cytochrome b5 reductase type 3, isoform 2; accession number NP_015565.1) and CYB5B (cytochrome b5 type b, accession number NP_085056.2) were expressed in *Escherichia coli* and purified as previously reported [22]. The expression plasmid for CYB5b (pET11a:CYB5), [34] encoding the soluble portion of the human mitochondrial CYB5, was a kind gift from Dr. Mario Rivera (University of Kansas}. Human CYB5R3 and CYB5b were expressed and purified as described previously [22]. CYB5R3 concentrations were calculated using an extinction coefficient of 10.6 mM^-1^cm^-1^ at 461nm [35]. CYB5b concentrations were determined using an extinction coefficient of 117 mM^-1^cm^-1^ at 413nm for the ferric form [36].

The DNA sequence encoding the human CYB5R4 protein was synthesized as a gBLOCK double stranded DNA fragment (IDT, Coralville, IA) with 3’and 5’ flanking regions complementary to the pET28 plasmid NdeI and HindIII restriction sites. The recombinant protein including an N-terminal Histag was expressed and purified similarly to the CYB5R3 protein [22]. CYB5R4 concentrations were calculated using an extinction coefficient of 185 mM^-1^cm^-1^ at 410 nm for the fully oxidized protein [32].

Protein purity was assessed by SDS-PAGE and UV−visible spectroscopy; spectral data was used to compare the protein properties to previously reported standards [7, 8, 16]. All proteins were buffer exchanged into 100 mM sodium phosphate, pH 7.4 (Fisher Scientific, Pear Lawn, NJ) and stored at −80 °C

### Kinetics of Globin reduction by CYB5R3

Reactions of Ngb/Cygb with the CYB5R3/CYB5B reducing system were conducted as previously described with minor modifications [22, 23]. The reaction was studied in pseudo first order conditions. Reaction mixtures included 1 nM CYB5R3, 10 μM Globin, and a variable amount of CYB5b (10 nM to 1.5 μM). The reactions were started by adding NADH (100 μM). Spectral changes were followed over time via UV-Visible spectrophotometry. Spectra were collected under anaerobic conditions in a glove box (Coy Labs, Grass Lake, MI) to prevent formation of the oxygen bound ferrous Cygb/Ngb upon reduction, resulting in convoluted spectral changes as well as autoxidation concerns. Reactions were studied at 37 °C either in 100 mM Sodium Phosphate, pH 7.4, or Phosphate buffered saline, pH 7.4.

### Kinetics of Globin reduction by CYB5R4

Reactions of Ngb/Cygb with CYB5R4 were studied in pseudo first order conditions. Reaction mixtures included a variable amount of CYB5R4 (100 nM to 1 μM) and 10 μM Globin. The reactions were started by adding NADH (100 μM). Spectral changes were followed over time via UV-Visible spectrophotometry. As in the case of CYB5R3, reactions were monitored under anaerobic conditions in a Glove box (Coy Labs, Grass Lake, MI). Spectral changes were fit to single exponentials to determined observed rate constants. Reactions were studied in Phosphate buffered saline, pH 7.4, at 37 °C.

### Neuroglobin/Cytoglobin NO dioxygenase activity

The reaction of the ferrous-oxy (Fe^2+^-O_2_) complexes of Ngb and Cygb with NO in the presence of CYB5R4 was studied as described for the CYB5R3/CYB5 system [22]. To determine the ability of the reducing system to maintain Cygb/Ngb ferrous oxy complex, we studied the reaction as follows: in a 1 mL cuvette, closed with a screw cap with a rubber septum, 20 μM ferric Ngb/Cygb was mixed with 750 nM CYB5R4 and 200 μM NADH in buffer saturated with 100% oxygen. The reaction was monitored by UV−vis spectroscopy until the spectra stabilized showing characteristic peaks for oxyCygb/oxyNgb. Then NO-saturated buffer was added to the cuvette to achieve a final NO concentration of ≈ 5 μM. The spectral changes were monitored until the spectra reached again a stable, oxyCygb/oxyNgb species and then NO was added again; this process was repeated for 3−5 cycles. The reaction was studied on an HP8453 UV−vis spectrophotometer (Agilent Technologies, Palo Alto, CA). Spectral deconvolution, based on Cygb/Ngb standard spectra was used to calculate the amount of each globin species (oxy, deoxy, met, ferrous-NO) during the reaction. The calculated concentrations of deoxyCygb and deoxyNgb during the reaction were negligible. Reactions were followed at 37 °C in phosphate buffered saline, pH 7.4.

### Modeling of Cygb-CYB5R4 and Ngb-CYB5R4 complexes

The computational models of the Cygb-CYB5R4 and Ngb-CYB5R4 complexes were generated by AlphaFold3 [37] or Chai-1 [38] using the sequences for wild-type human Cygb/Ngb and the *b*_5_ domain sequence of human CYB5R4 (residues 51-137). Two heme ligands were included in the modeling to account for both globin and *b*_5_ domain heme moieties. The heme domains were properly placed by the models in the known heme pockets. No restrictions in terms of interface residues were used. Buried surface was calculated as the difference between the unliganded and liganded structures for each model; individual surface areas were determined with the Surface Racer software, v5.0 [39] using a 1.4 Å probe radius. The interaction surface was calculated as the sum of the buried surfaces for both proteins divided by 2. Figures were made using PyMOL, version 0.99rc6 [40].

## RESULTS

### Reevaluation of Cygb/Ngb reduction by NADH/CYB5R3

We have previously shown that Cygb is reduced by the CYB5R3/CYB5 system [22, 23]. Although we did not observe significant reduction of Ngb, we note that the experiments were conducted in supraphysiological ionic strength conditions (100 mM Sodium phosphate buffer, pH 7.4). Considering that the interactions between CYB5 and its electron acceptors are highly electrostatic in nature [22, 41, 42] the reaction between CYB5 and Ngb at physiological ionic strength may occur at a faster rate compatible with physiological function. To verify this point we determined the reduction rates for Cygb and Ngb in physiological ionic strength conditions using PBS.

In our previous work, we have determined the reduction rate of Cygb by the CYB5R3 system [22] with a rate constant for electron transfer (*k_et_*) from the electron mediator hemoprotein cytochrome *b_5_*b (CYB5b) to the terminal hCygb of 2.87 × 10^5^ M^-1^s^-1^ at pH 7.4 and 37°C. We did not detect significant electron transfer to Ngb [22]. Here we reexamined the reaction in PBS to assess whether the ionic strength of the solution influences the reaction rate. In a quartz cuvette we mixed 1 nM CYB5R3, 10 µM WT Cygb, and a variable amount of CYB5b (10 nM to 250 nM). 100 µM NADH was then added to initiate the reaction. The observed rates are shown in Figure 2. A linear fit of the values yields a bimolecular reaction rate for Cygb of 4.19 × 10^6^ M^-1^s^-1^ at pH 7.4 and 37°C; about 10-fold faster than the observed reaction in 100 mM sodium phosphate [22].

**Figure 2.**
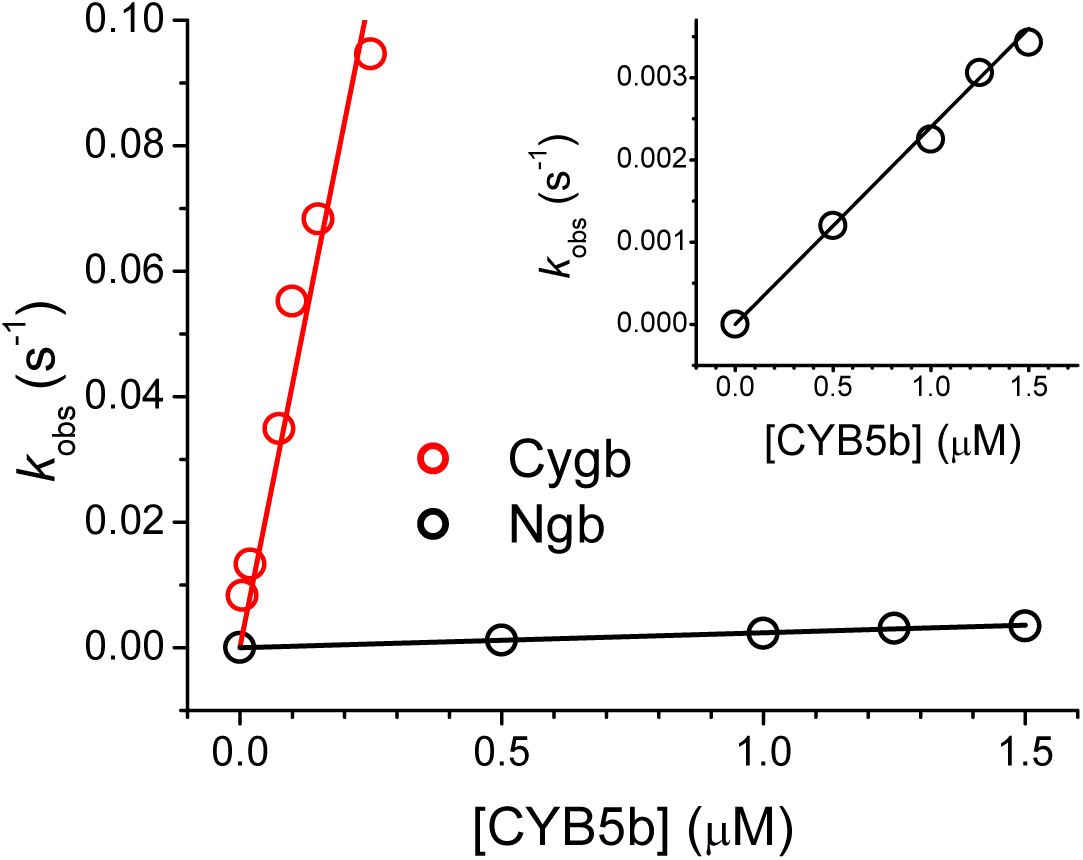
Cygb is rapidly reduced by the NADH/CYB5R3/CYB5B system, but not Ngb. The reactions are studied under pseudo first order conditions (50 nM CYB5R3, 5-15 µM WT Cygb/Ngb, and a variable amount of CYB5b (100 to 250 nM for Cygb, 500 nM to 1.5 µM for Ngb). The reduction was initiated by the addition of 100 µM NADH. Reactions were carried out in phosphate buffered saline at pH 7.4 and 37°C. The inset shows the neuroglobin data independently.

In a manner analogous to the Cygb determination under pseudo first order conditions, 1 nM CYB5R3, 10 µM WT Ngb, and a variable amount of CYB5b (500 nM to 1.5 µM) were added to a cuvette. 100 µM NADH was then added to initiate the reduction; the spectral changes were followed over time via UV-Visible spectrophotometry. Moreover, spectra were collected under a nitrogen atmosphere to prevent formation of the oxygen bound ferrous Ngb upon reduction, resulting in convoluted spectral changes as well as autoxidation concerns. Spectral changes over time at 559 nm, the main Q-band peak of unliganded reduced Ngb (deoxyNgb), were fit to single exponentials to determined observed rate constants, and the second order electron transfer rate constant of 2.40 x 10^3^ M^-1^s^-1^ was derived from standard kinetic techniques via changing the concentration of CYB5b (Figure 2; Supplemental Figure 2). Controls with CYB5R3, NADH, and either Ngb or Cygb indicate that the flavin containing protein is only capable of slowly reducing Cygb, and shows no reduction of Ngb (data not shown)[22]. This is consistent with other observations indicating a lack of direct reduction of Cygb by CYB5R3 [43]. Such a small rate constant of electron transfer from CYB5b to Ngb and relative local concentrations suggests that this reducing system would be unable to maintain the reduced ferrous Ngb population needed in vivo for its proposed functions. Thus, we conclude that human Ngb is not efficiently reduced by the NADH/CYB5R3 system

### Human Cygb and Ngb are both readily reduced by the NADH/CYB5R4 system

Next, we investigated the ability of CYB5R4 to reduce Cygb and Ngb. Unlike CYB5R3, in CYB5R4 a CYB_5_b-type cytochrome is fused to the flavin reductase domain, eliminating the need of a partner electron transfer protein before interaction with the terminal hemoprotein. Reduction of the flavin subunit by NADH is much faster than the subsequent electron transfer steps from the flavin to the CYB5 moiety and then from the CYB5 heme to the terminal electron acceptor, analogous to the NADH/CYB5R3/CYB5b system. Thus, varying the concentrations of CYB5R4 (100 nM-1 µM) allows for determination of the second order electron transfer rate constants (*k_et_*) from CYB5R4 to the terminal hemoprotein of interest. We monitored the reduction of 10-20 µM of the target hemoprotein by CYB5R4, with the reaction initiated by the addition of 100 µM NADH. Experiments were performed under an oxygen-free atmosphere to remove autoxidation as a factor as well as limit the observed reduced species to only the reduced deoxy-species.

The observed rates for the reaction of CYB5R4 and selected heme proteins are shown in Table 1. We observe that like CYB5R3, wild-type human Cygb is rapidly reduced by CYB5R4 (see Figure 3A, Table 1). Notably, we observe that unlike CYB5R3, the CYB5R4 readily reduces wild-type human Ngb (Figure 3B). Figure 3B depicts 167 nM CYB5R4 added to 15 µM ferric WT Ngb, initiated with 100 µM NADH, and formation of the signature hexacoordinate deoxyNgb Q-bands. Following the formation of deoxyNgb at 563 nm and the decay of ferric Ngb at 497 nm over time after reaction initiation and fit to a single exponential gives the *k_obs_* at a given concentration of CYB5R4 (Figure 3E). The second order *k_et_* was determined to be 5.13 × 10^4^ M^-1^s^-1^ (Table 1, Figure 3F), over 20-fold higher than the rate previously determined for the CYB5R3/CYB5b couple (2.40 × 10^3^ M^-1^s^-1^).

**Figure 3.**
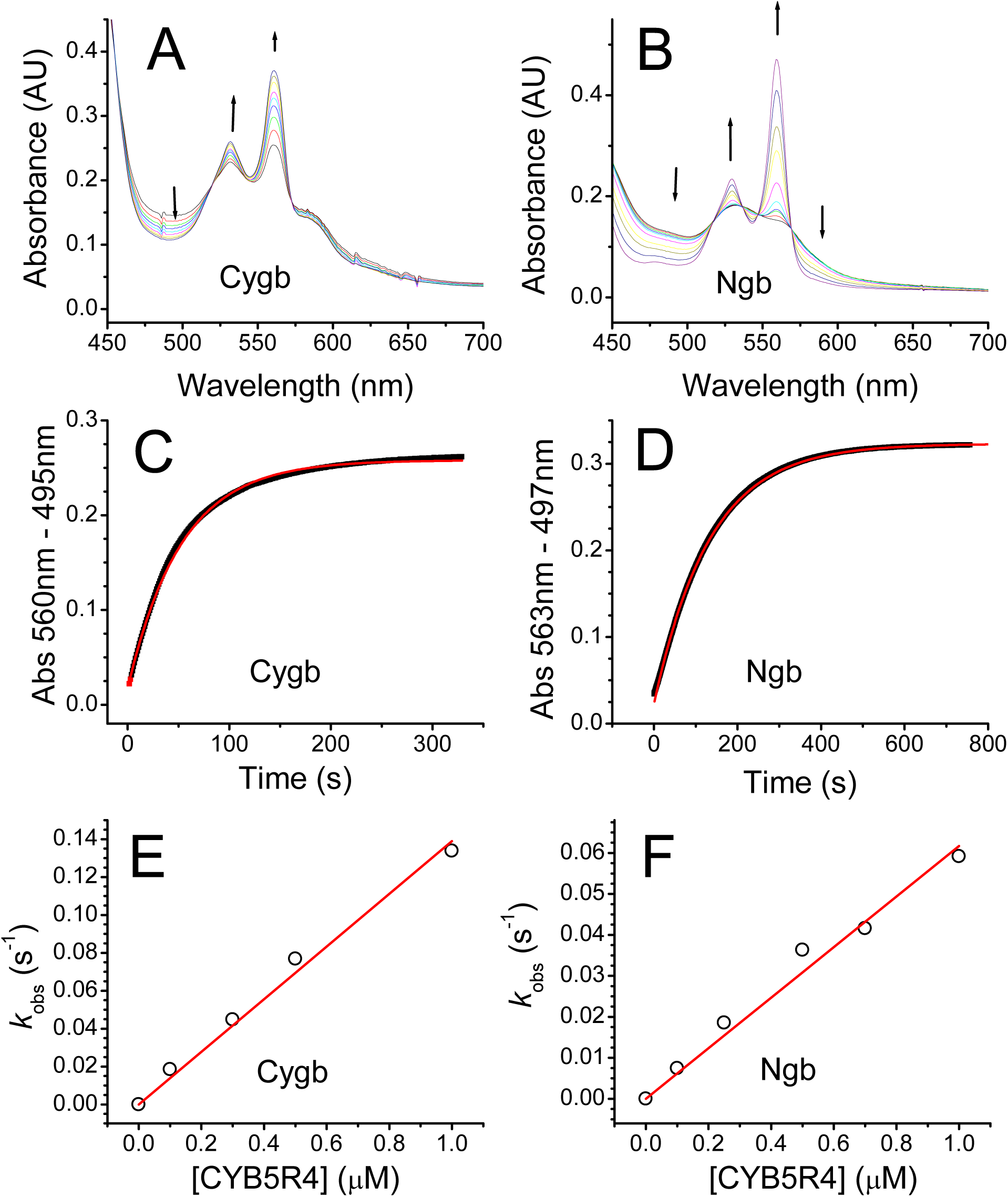
Determination of reaction rate constants for the reaction of CYB5R4 with Cygb/Ngb. Panels A-B, spectral changes during the reaction of 100 nM CYB5R4 with wild type Cygb (A) or wild-type Ngb (B) in the presence of 100μM NADH. Panels C-D, absorbance changes at 560nm-495nm (WT Cygb, Panel C) or 563nm-497nm (WT Ngb, Panel D) during the reaction of CYB5R4 with each globin as depicted in Panels A and B, respectively. Absorbance changes are fitted to a single exponential equation to determine observed rates. Panels E-F, observed rates as a function of the concentration of CYB5R4 used. A linear fit of the observed rates yields a slope equal to the bimolecular rate constant for the reaction.

**Table 1.** Second order electron transfer rate constants between CYB5R4 and different heme proteins. Mutations in Cygb and Ngb change interaction of *b*_5_ domain with the terminal hemoprotein in PBS at 37°C.

| Protein | $k_{\text{et}} (\text{M}^{-1}\text{s}^{-1})$ |
| --- | --- |
| *Hemoglobin* |  |
| WT | $1.05 \times 10^4$ |
| *Myoglobin* |  |
| WT | $3.22 \times 10^4$ |
| *Neuroglobin* |  |
| WT | $6.17 \times 10^4$ |
| D73K | $5.43 \times 10^4$ |
| E60K | $7.68 \times 10^4$ |
| E87K | $5.62 \times 10^4$ |
| *Cytoglobin* |  |
| WT | $1.54 \times 10^5$ |
| C38S/C83S | $4.84 \times 10^5$ |
| L46H | $2.90 \times 10^4$ |
| H81A | $8.39 \times 10^4$ |
| H81Q | $7.15 \times 10^4$ |
| V85I | $1.98 \times 10^5$ |
| R84E | $4.27 \times 10^4$ |
| K116E | $3.52 \times 10^4$ |

### CYB5R4 reduces preferentially Cygb and Ngb over other human globins

Like the CYB5R3/CYB5 system, CYB5R4 is expected to catalytically reduce hemoproteins or other one-electron acceptors using NAD(P)H as a reductant. However, the physiological targets of CYB5R4 are unknown [44]. In order to determine if known mammalian globins are possible physiological targets of CYB5R4, we studied the reduction of these proteins (Hb, Mb, Cygb, Ngb) by CYB5R4. Except for WT Cygb (Figure 4), the reduction rate of Ngb by 500 nM CYB5R4 (Figure 4) is faster than other mammalian globins. Hemoglobin (Hb, Figure 4) and myoglobin (Mb, Figure 4) are both reduced nearly an order of magnitude slower under identical conditions (Table 1, Supplemental Figure 3). Altogether, CYB5R4 is more efficacious towards the reduction of Cygb/Ngb than towards Hb/Mb.

**Figure 4.**
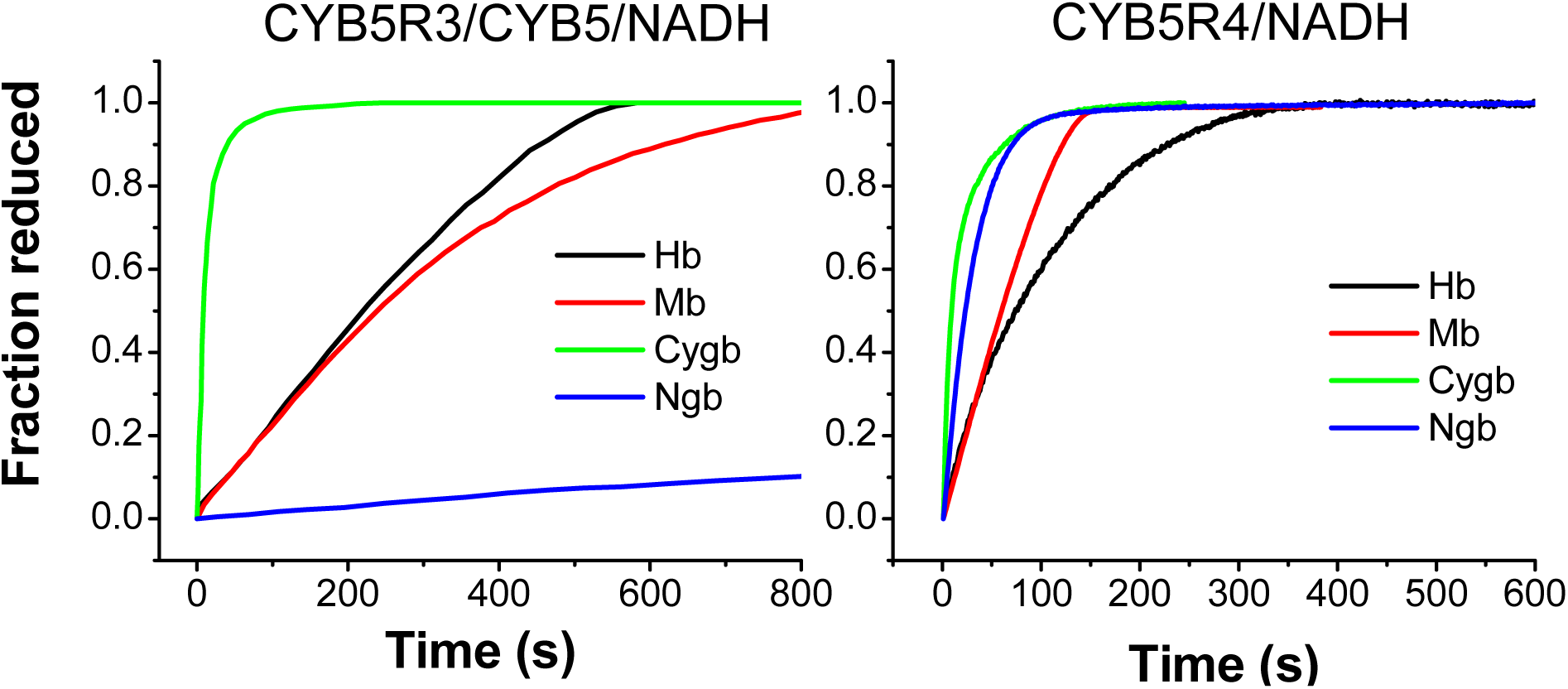
Reduction of globins by the CYB5R4 system as compared to the CYB5/CYB5R3/NADH. Left, reduction of ferric wild-type globins by the CYB5R3/CYB5/NADH system. Globins (≈20 μM) were incubated with 2 μM human CYB5b and 0.2 μM human CYB5R3. The reaction was initiated by the addition of 100 μM NADH. All experiments were performed under anerobic conditions, in 100 mM phosphate buffer, pH 7.4, at 37°C. Data from ref [22]. Right, reduction of different ferric wild-type globins by 500 nM CYB5R4 in the presence of 100 μM NADH. Reactions were performed under anerobic conditions, in phosphate buffered saline pH 7.4, at 37°C.

### Effect of mutations on the interaction of Ngb and Cygb with CYB5R4

Once we established the reducing capacity of CYB5R4 on Ngb and Cygb, we investigated the effect of several mutations on the reduction rates to help determine the nature the electron transfer from CYB5R4 in Cygb and Ngb. Most mutations that were probed investigated the probable interface between the two proteins, where ionic interactions are expected to facilitate electron transfers between the two hemes [22, 36, 37]. The mutations studied for both Cygb and Ngb and the observed rate constants are listed in Table 1.

Cygb mutations L46H, H81A, H81Q, and V85I are located within the distal heme pocket, and although they modify the heme properties to different extent, they are not solvent exposed and would be unable to directly interact with CYB5R4. On the other hand, C38S/C83S, R84E and K116E mutations involve surface residues that could be involved in the binding of CYB5R4.

Formation of the disulfide between Cys38 and Cys83 weakens the interaction between the heme iron and the distal histidine (His81), causing some opening of the heme cavity; a similar effect is caused by His81 mutations [10, 15, 16]. The observed rates for the reduction of the mutants by CYB5R4 indicate that a more open heme pocket leads to slower reduction rate that that observed for the WT protein (e.g. H81A and H81Q, Table 1, Supplemental Figure 4) but the C38S/C83S mutant, which cannot form the disulfide bond and is locked in a more “closed” heme pocket conformation, shows faster reduction rates (Table 1, Supplemental Figure 5).

Residues Arg84 and Lys116 flank the outer part of the heme pocket and coordinate the propionate groups of the heme (Figure 5). These residues have been postulated as mediators of ascorbate binding to Cygb [21]. They are also important for lipid binding to Cygb [10]. We observe that the introduction of negative charges in these positions lead to a 4-fold decrease in reduction rates (Table 1, Supplemental Figure 5), indicating that the binding of CYB5R4 to Cygb occurs, at least in part, around the positively charged region outside the heme pocket [10] (Figure 5).

**Figure 5:**
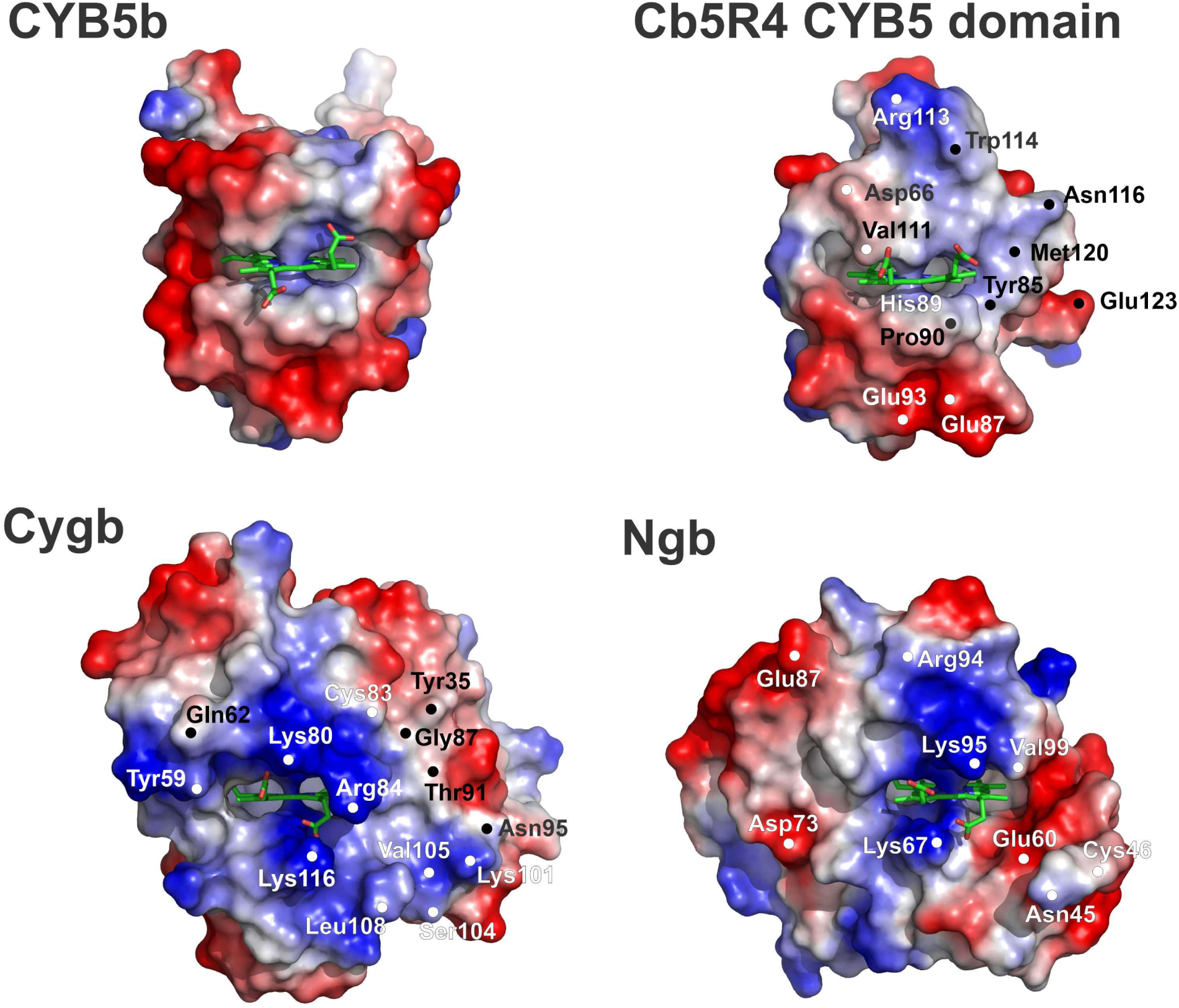
Electrostatic surfaces of Ngb, Cygb, CYB5b and the B5 moiety of CYB5R4. The surface potentials for human CYB5b, the cytochrome *b*_5_ domain of CYB5R4, human Cygb and human Ngb are shown. Positive potential is represented in blue, negative surface potential in red. The heme group is represented as green sticks. The PDB structures used are 3NER (CYB5), 3LF5 (CYB5R4 cytochrome *b*_5_ domain), 1UMO (hCygb), and 1OJ6 (Ngb). Selected surface amino acids around the exposed heme and/or identified in the docking interfaces are marked. Protein structures and electrostatic surface potentials were generated with PyMOL [39].

Ngb mutations E60K, D73K and E87K are surface mutations previously studied in the context of the Ngb-Cytochrome *c* interaction[33]. We do not observe substantial changes in the bimolecular rate constant for Ngb reduction, except for a 1.5-fold increase in the case of the E60K mutant (Table 1, Supplemental Figure 6). These results suggest that Asp73 and Glu87 are not in the binding interface, whereas Glu60 is located in the interface and probably in proximity of some negatively charged patch in the CYB5R4 surface. Together with the Cygb results, the data suggest that negatively charged patches in the CYB5R4 surface mediate the interaction with both globins.

For electron transfer to readily occur, the two heme pockets must be relatively close; efficient transfer is thus facilitated by appropriately matched electrostatics. As CYB5R4 appears to have a surface potential that is predominantly negative (Figure 5), mutations of positive residues Arg84 and Lys116 to acidic residues in Cygb should result in slower electron transfer kinetics, whereas mutation of Glu60 in Ngb to a basic residue should increase electron transfer kinetics. The corresponding *k_et_* for mutant and WT proteins are shown in Table 1.

### Docking studies of the Cygb/CYB5R4 and Ngb/Cyb5R4 interaction

In order to get a clearer picture of the possible interactions between Cygb/Ngb and CYB5R4, and with the additional guidance of the empirical data reported above, we investigated the protein-protein interaction using computational methods. To assess the potential structures of the complex we used AlphaFold3 [37] and Chai-1 [38] using the sequences for wild-type human Cygb/Ngb and the *b*_5_ domain sequence of human CYB5R4 (residues 51-137).

We did not impose initial constraints to the calculations, letting the docking algorithms to explore the whole protein surfaces. To our surprise, even without restrictions, most of the models showed the globins and the CYB5R4 heme domain interacting through their sides where the heme is solvent exposed (Figure 6). Fast electron transfer depends on different factors, but the distance between redox centers is critical, with most biological interactions occurring within distances of 14Å or less [45]. Thus, the resulting complexes were filtered by the shortest heme-heme distance. Out of 10 models per complex, 5 Cygb/CYB5R4 models and 7 Ngb/CYB5R4 models placed the heme groups in proximity (≤6 Å). Selected models depicting possible interaction structures for Cygb/CYB5R4 and Ngb/CYB5R4 are shown in Figure 6 and Supplemental Figures 7, 8 and 9. The models with protein contacts more consistent with the experimental data (Figure 6C and D; Supplemental Figure 9) correspond to the Chai-1 model 1 for both Ngb and Cygb; further model details are presented in the Supplemental Table 1 and Supplemental Figures 7, 8 and 9.

**Figure 6.**
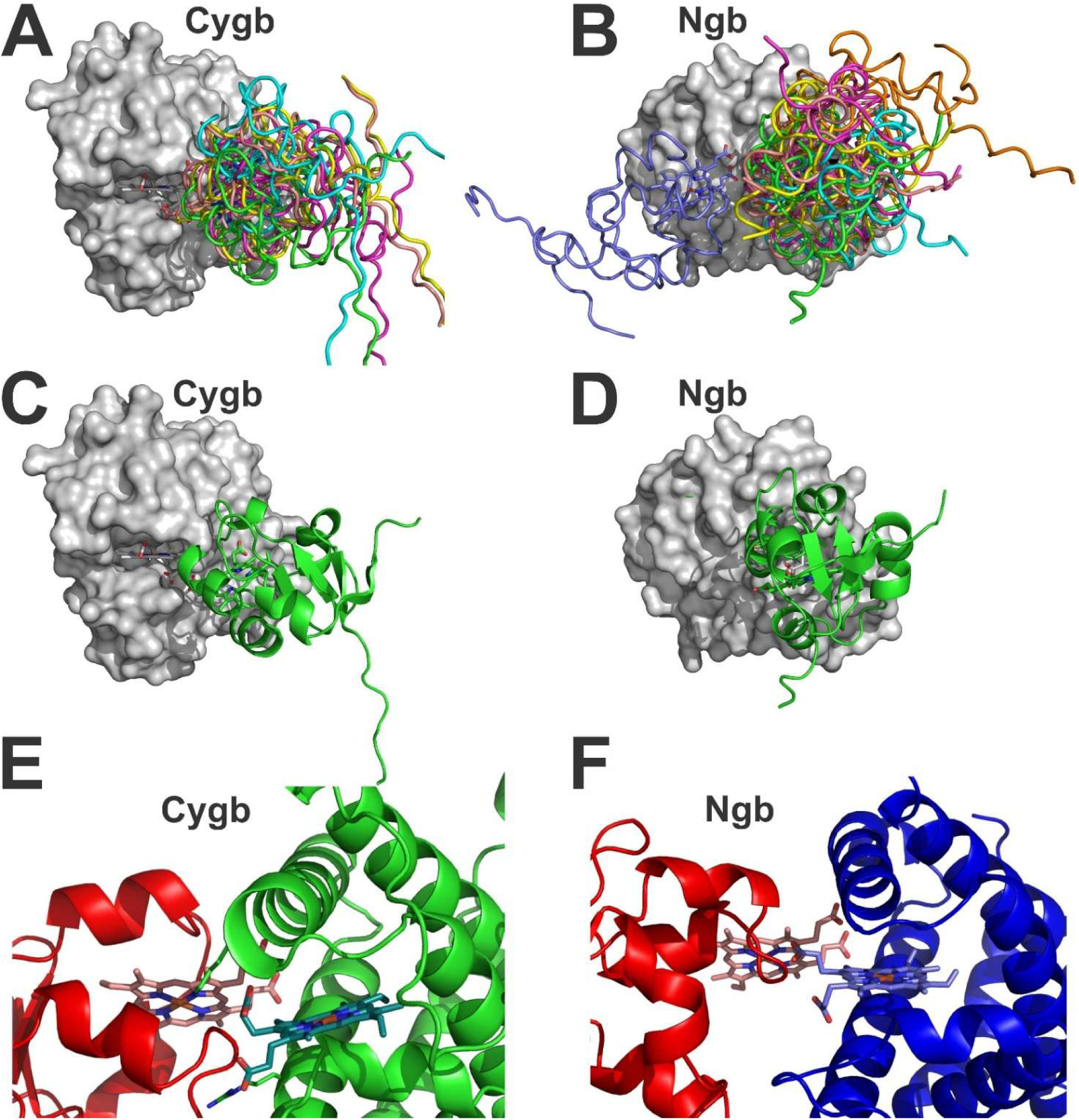
Docking models for the interaction of the cytochrome *b*_5_ domain of CYB5R4 with Cytoglobin and Neuroglobin. Panels A and B show the 5 selected Cygb/CYB5R4 complexes (Panel A) and the 7 selected Ngb/CYB5R4 complexes (Panel B). The noodle models show the CYB5R4 location in each model. Cygb and Ngb molecular surfaces are shown in similar orientation to that in Figure 5, with the heme moiety depicted as sticks. Panels C and D show the model structure most consistent with the experimental data (Chai-1 model #1) for Cygb (Panel C) and Ngb (Panel D). A more detailed view of the heme interactions is shown in Panels E (Cygb) and F (Ngb). The CYB5R4 cytochrome *b*_5_ domain (red) is shown in a similar orientation in both complexes. Cygb (green) and Ngb (blue) appear to bind in a slightly different orientation, although the heme-to-heme distances are adequate for fast electron transfer (<6Å) in both complexes. Figure generated with PyMOL [39].

Most of the resulting models showed consistent features with the experimental results. For the Cygb/CYB5R4 models, all the selected models included Cygb Arg84 in the interaction surface, and half of them also included Cygb Lys116 in the surface. These models are considered more consistent with the experimental results and thus more plausible (Figure 6C-F, Table 1, Supplemental Table 1, Supplemental Figures 7 and 9).

For the Ngb/CYB5R4 interaction, Ngb residues Asp73 and Glu87 are largely absent from the calculated interfaces, in agreement with our results (Figure 6B, Table 1, Supplemental Table 1, Supplemental Figures 8 and 9). There is only one model showing E60 in the interface. The mutation of Glu60 to Lysine in this model could allow for interactions with the main chain of the CYB5R4 Val111 residue or the heme carboxylate, but may also allow to avoid the formation of non-productive complexes, so the selective value of this contact is unclear. Altogether, the interaction surface for Ngb and CYB5R4 seems to be much smaller than the area used when Ngb binds to cytochrome *c* [33].

### In vitro assessment of redox cycling based of NO dioxygenase activity

Ferrous oxygen-bound hemoproteins rapidly react with NO to yield ferric hemoprotein and nitrate in a process called NO dioxygenation (Eq. 1):

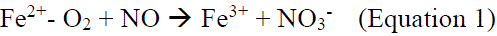

Most oxygen-bound hemoproteins react with NO at a rate constant greater than 10^7^ M^-1^s^-1^, up to 2.9×10^9^ M^-1^s^-1^ in some flavohemoglobins [46], well beyond the rate-limiting rate constant of electron transfer from CYB5R4 to a terminal hemoprotein measured in our work (Table 1). In this context, we have shown that NO dioxygenation can be used as a proxy to evaluate the catalytic efficiency of a hemoprotein reducing system such as CYB5R3/CYB5 or CYB5R4 in physiological-like conditions [22]. In our assay, the globin is mixed with the reducing system and NADH is added to initiate protein reduction and generate a fully oxygenated species. Once the 100% oxy species is formed, NO –about half of the heme protein concentration-is injected in the cuvette causing a sudden decay in the ferrous deoxy species and simultaneous increase in the ferric species. The ferric species is reduced back to ferrous-oxy species by CYB5R4 according to the specific rate for each heme protein. Once the ferrous oxy species is fully regenerated, NO can be injected again, and the process is repeated. The rate of ferrous oxy decay upon NO addition is fast and largely protein independent; the rate of ferrous oxy formation once all NO is consumed is directly related to the specific reaction rate between the reductant and the heme protein. Here we investigated the NO dioxygenase activity of WT Ngb and Cygb with CYB5R4, as well as the reaction for the mutants that showed a greater change in reductase activity vs WT, Cygb R84E and Ngb E60K (Table 1)

The reaction for WT Cygb and R84E Cygb is shown in Figure 7 (top panels). Addition of NO causes a quick formation of ferric heme, but the CYB5R4 is able to fully reduce the protein back to the ferrous-oxy species in <50s. However, in the case of R84E Cygb, the reduction process is severely impaired; the full regeneration of the ferrous-oxy species takes >200s. In addition, the steady state level of ferrous oxy protein does not reach 100%, an observation that we have also noted when ascorbate is used as a reductant for wt Cygb [22]. This suggests that the reduction rate is not fast enough to fully outcompete protein autoxidation. Similar results are observed with the K116E mutant.

**Figure 7:**
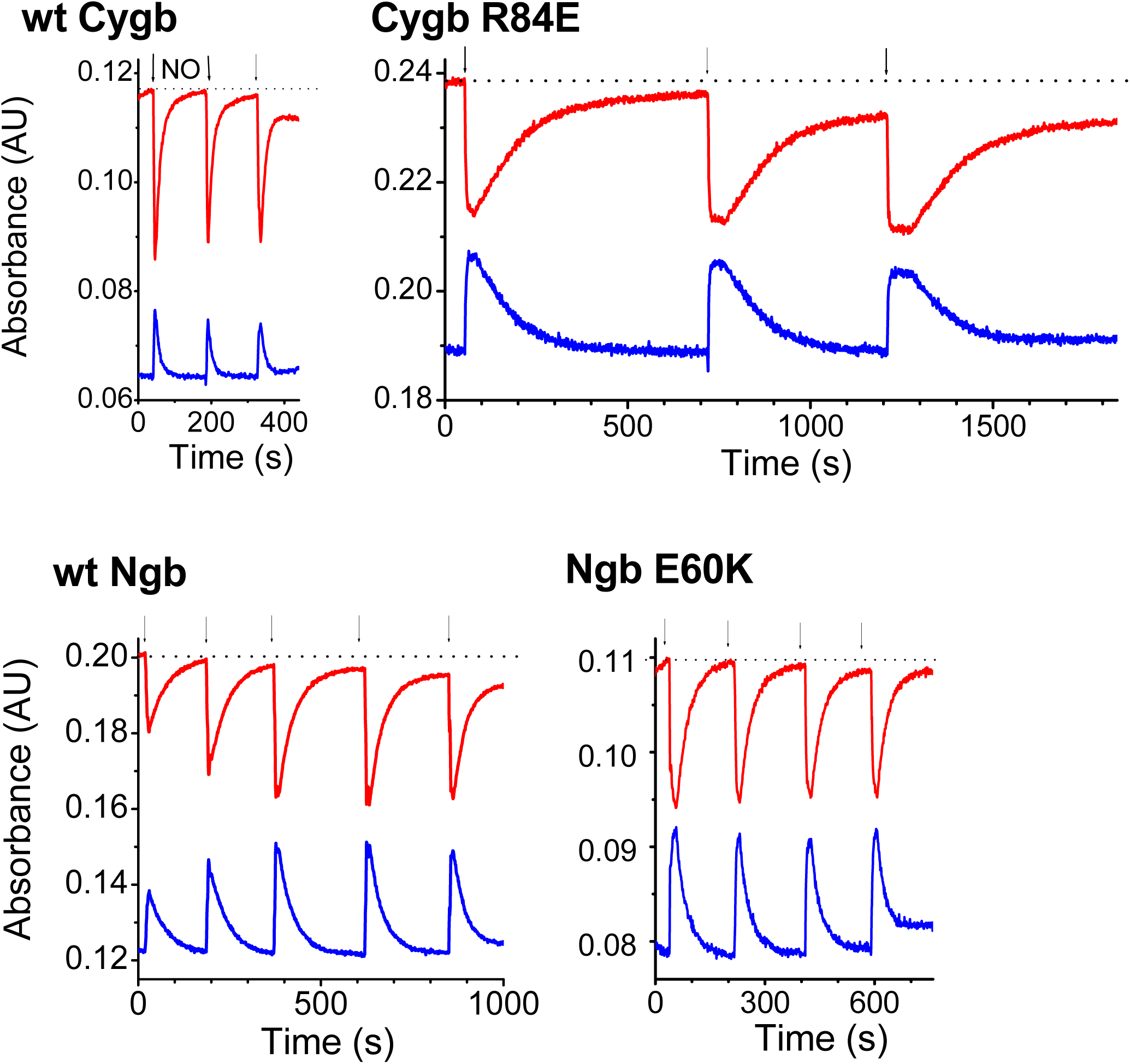
NO dioxygenation by Ngb, Cygb, and select mutants in the presence of CYB5R4 and NADH. Globin samples in aerobic buffer are in a mixed ferric/ferrous oxy state which decays slowly to a fully ferric species in the absence of reductant. In the presence of 750 nM CYB5R4, upon addition of 200 μM NADH in O_2_-saturated buffer, ferric globin is reduced (blue lines, 521nm for Cygb, 525nm for Ngb) with concomitant formation of ferrous-oxy globin (red lines, 542nm for both Cygb and Ngb). After 100% oxy species is formed, NO (about half of the protein concentration) is injected in the cuvette causing a sudden decay in the ferrous deoxy species and simultaneous increase in the ferric species. The ferric species is reduced back by CYB5R4 according to the specific rate for each protein. The scale in the x-axis is similar in all panels to allow for direct comparison. Arrows indicate the additions of nitric oxide.

The reaction for WT Ngb and E60K Ngb is shown in Figure 7 (bottom panels). It can be noted that the reduction of Ngb is fast (around 150 s) but about half the rate of the reduction of Cygb, consistent with the observed direct reduction rates (Table 1). Notwithstanding, the CYB5R4 reducing activity is able to maintain the protein mostly in the ferrous-oxy state and represents, to our knowledge, the first reductant able to efficiently maintain a stable ferrous Ngb population in vitro. There is some decay on the final ferrous-oxygen levels after each NO addition, but not as marked as for the Cygb R84E mutant. We also investigated the reaction for the E60K, as it showed a faster reduction rate in the direct reduction assay (Table 1). Consistent with our previous observations, the mutant is reduced faster than wt Ngb (around 100s) but slower than wt Cygb. CYB5R4 maintains ferrous oxy levels around 100% after repeated NO additions, indicating that the reduction rate outpaces the heme autoxidation rate.

## Discussion

The affinity of ferrous heme proteins towards ubiquitous oxygen, and the intrinsic oxidation rates of these ferrous-oxy complexes, makes necessary the presence of cellular systems capable of fast heme reduction. In the case of hemoglobin, CYB5R3 is well known as a key component of the reduction system; mutations in the soluble isoform of CYB5R3 are related to congenital methemoglobinemia [47]. As novel members of the globin family emerged, the nature of their reducing systems became an open question [26–28]. Cygb, Ngb and Androglobin are particularly relevant in that regard as they are found in the human genome and may be involved in processes of clinical relevance [1, 2].

Halligan et al. discovered the ability of Cygb to catalyze NO oxidation in vivo [48]; a process that necessarily requires a reduction system. Gardner et al. were the first to show the ability of several systems to reduce Cygb in vitro and support NO dioxygenation reactions, including ascorbate, CYB5R3, CYB5R4 and cytochrome P450 reductase[21]; however, they concluded that in their experimental conditions none of the electron donors was capable of supporting NO scavenging activity for Ngb (or Mb) [21]. Studies by Ilangovan et al. demonstrated that in vascular smooth muscle cells CYB5R3/CYB5 appear to support >80% of Cygb NO dioxygenase activity, but some residual activity is independent of CYB5R3/CYB5 [19]

Here, we demonstrate that CYB5R4 can reduce both Cygb and Ngb with a high reduction rate compatible with a physiological function, that could be related, but not necessarily limited to NO scavenging. Other possible functions, such as oxygen transport or nitrite reduction would also require efficient reduction systems [7]. This is the first efficient reductant identified for Ngb. Jusman et al. have investigated the ability of bovine brain and liver extracts to reduce Cygb and Ngb [25, 49] and found that extracts around 60kDa had Cygb reductase activity, albeit weak [49]. Similarly, a fraction including a band between 50 and 75kDa showed some reductase activity against Ngb [25]. It is conceivable that both fractions may include CYB5R4 (MW 59.5 kDa). Although CYB5R3 can also reduce Cygb, its activity without CYB5 is weak [22], and may be hard to identify following size-exclusion fractionation. However, we consider that these reports are consistent with CYB5R4 being a main reductant of Cygb/Ngb in vivo.

We also present docking complexes for the Cygb/CB5R4 and Ngb/CYB5R4 interaction that aer in good agreement with the experimental data. The surface areas for the complexes of Cygb or Ngb with the *b*_5_ domain of CYB5R4 were around 300-600 Å^2^, in the lower end of the areas observed for most protein-protein complexes (usually in the 500-2000 Å^2^ range, [50]). This is also consistent with previous observations indicating that electron transfer complexes often use small surface areas balancing binding with fast turnover rates [51]. The surface area for Cygb was larger in average (460±110 Å^2^ for Cygb/CYB5R4 vs 360±90 Å^2^ for Ngb/CYB5R4). Interestingly, in both complexes (Cygb/CYB5R4 and Ngb/CYB5R4) one of the globin disulfide bond-forming cysteines (Cys83 in Cygb, Cys 46 in Ngb) seems consistently involved on the binding interface; this suggests that some interplay between disulfide bond formation and interaction with CYB5R4 is possible. It is also intriguing considering that some asymmetry has been observed in Cygb thiol reactivity, which points out roles for these cysteine residues beyond the thiol/disulfide equilibrium. Recently, Jourd’heuil et al. have reported that mutation of Cys83 abolishes Cygb hydrogen peroxide reactivity, whereas mutation of Cys38 shows no effect [6]. Whether similar pathways operate for Ngb and are directly relying on interactions with CYB5R4 is an intriguing hypothesis.

The strong reducing effect of CYB5R4 on Ngb may have translational impact. A specific engineered mutant of Ngb – Ngb with an H64Q substitution combined with three surface thiol substitutions (C46G/C55S/C120S; Ngb-H64Q-CCC) – has been demonstrated to serve as a potential carbon monoxide (CO) scavenging agent [52, 53]. CO poisoning is a condition that impacts over 50,000 persons in the United States annually [54, 55]. These agents have also been postulated as artificial oxygen carriers [56] and potential blood substitutes. Again, a major technical challenge for the use of such molecules as oxygen carriers could be maintaining Ngb, Cygb or other heme proteins in their reduced, or active state. Ngb-H64Q-CCC has an autoxidation rate of 0.86 hour^−1^, which is not an obstacle for fast CO binding when infused in ferrous form, but is not optimal for oxygen carrier functions [53]. Unlocking an enzymatic highly efficient reducing system could enable the further development of these agents as CO poisoning antidotes or blood substitutes.

Analysis of CYB5R3 and CYB5R4 expression in mammalian tissues [57, 58] indicates that both proteins are ubiquitously expressed and also found in the cytoplasm [44] as observed for Ngb/Cygb (although nuclear localization of Cygb/Ngb has been also reported [13, 59]); it is thus expected that they will be able to mediate Cygb/Ngb reduction. Certainly, CYB5R3 and CYB5R4 are involved in other known and unknown cellular roles. CYB5R3 is present in most cells where it participates in lipid metabolism, heme protein reduction and mediates oxidative responses [29]. CYB5R4 was discovered merely 25 years ago, and its functions are not fully elucidated, although by similarity to CYB5R3 a role in lipid metabolism was postulated [31]. The complete deletion of the CYB5R4 gene in mice causes progressive loss of pancreatic beta cells, apparently due to inability to cope with oxidative stress [60]; further studies of the CYB5R4 KO phenotype also found increased fatty acid catabolism and oxidative stress in liver cells [61]. Interestingly, early studies found Ngb in pancreatic islets of Langerhans [62]; overexpression of Ngb [63] was shown to improve viability of islet cells for transplantation purposes. Cytoglobin has been long-related to the oxidative stress response in liver [64, 65]. We speculate that both responses can be at least partly mediated by CYB5R4 function. These relationships may uncover novel pathways regulating the response to oxidants in vivo and deserve further investigation.

## Acknowledgements

We thank Mark T. Gladwin for funding support, encouragement and helpful discussions. O. C. was part of the Pittsburgh Undergraduate Research Diversity Program (PURDiP) at the Vascular Medicine Institute, University of Pittsburgh. E.D, S.K., S.M., M.H. and D.C. were part of the First Experiences in Research program at the University of Pittsburgh, Dietrich School of Arts and Sciences. We also want to acknowledge Lawrence Ji (First Experiences in Research program at the University of Pittsburgh, Dietrich School of Arts and Sciences) and Divya Jyoti and Jose Gutierrez (Summer Research Experience for High School Students program, University of Pittsburgh) for their work on this project. We also thank Kaitlin Bocian for excellent technical support.

## Abbreviations

CYB5R: cytochrome *b*_5_ reductase
CYB5R3: cytochrome *b*_5_ reductase isoform 3
CYB5R4: cytochrome *b*_5_ reductase isoform 4
CYB5a: cytochrome *b*_5_ type a
CYB5b: cytochrome *b*_5_ type b
Cygb: Cytoglobin
FAD: Flavin adenine dinucleotide
Hb: hemoglobin
Mb: myoglobin
NADH: nicotine adenine dinucleotide
NADPH: nicotine adenine dinucleotide phosphate
Ngb: neuroglobin

