## Supplementary data for "Cytochrome *b*_5_ reductase 4 efficiently reduces Neuroglobin and Cytoglobin"

*From the [1]Heart, Lung, Blood and Vascular Medicine Institute, University of Pittsburgh, Pittsburgh, PA 15261, [2]Division of Classical Hematology, University of Pittsburgh, Pittsburgh, PA 15261, [3]Department of Medicine, University of Maryland School of Medicine, Baltimore, MD 21201, USA, [4]Division of Pulmonary, Allergy and Critical Care Medicine, University of Pittsburgh, Pittsburgh, PA 15261, and the [5]Department of Bioengineering, Swanson School of Engineering, University of Pittsburgh, Pittsburgh, PA 15260 [6]Department of Pharmacology and Chemical Biology, University of Pittsburgh, Pittsburgh, Pennsylvania 15261, United States.*

### TABLE OF CONTENTS

**Supplementary Figure 1.** Domain arrangement and phylogeny of cytochrome b5 reductases.

**Supplementary Figure 2.** Spectral changes and reaction kinetics for the reaction of CYB5R3/CYB5b with wild type cytoglobin and neuroglobin in phosphate buffered saline.

**Supplementary Figure 3**. Spectral changes and reaction kinetics for the reaction of CYB5R4 with hemoglobin and myoglobin

**Supplementary Figure 4.** Spectral changes and reaction kinetics for the reaction of CYB5R4 with cytoglobin mutants L46H, H81A, H81Q, and V85I.

**Supplementary Figure 5.** Spectral changes and reaction kinetics for the reaction of CYB5R4 with cytoglobin mutants C38S/C83S, R84E, and K116E

**Supplementary Figure 6.** Spectral changes and reaction kinetics for the reaction of CYB5R4 with neuroglobin mutants E60K, D73K, and E87K.

**Supplementary Figure 7.** Modeled structures for the CYB5R4-Cytoglobin complexes using Alphafold3 and Chai-1

**Supplementary Figure 8.** Modeled structures for the CYB5R4-Neuroglobin complexes using Alphafold3 and Chai-1.

**Supplementary Figure 9.** Comparison of selected models for the interaction of the cytochrome $b_5$ domain of CYB5R4 with Cytoglobin and Neuroglobin

**Supplementary Table 1.** Main properties and interface residues for the selected CYB5R4/Cygb and CYB5R4/Ngb complexes

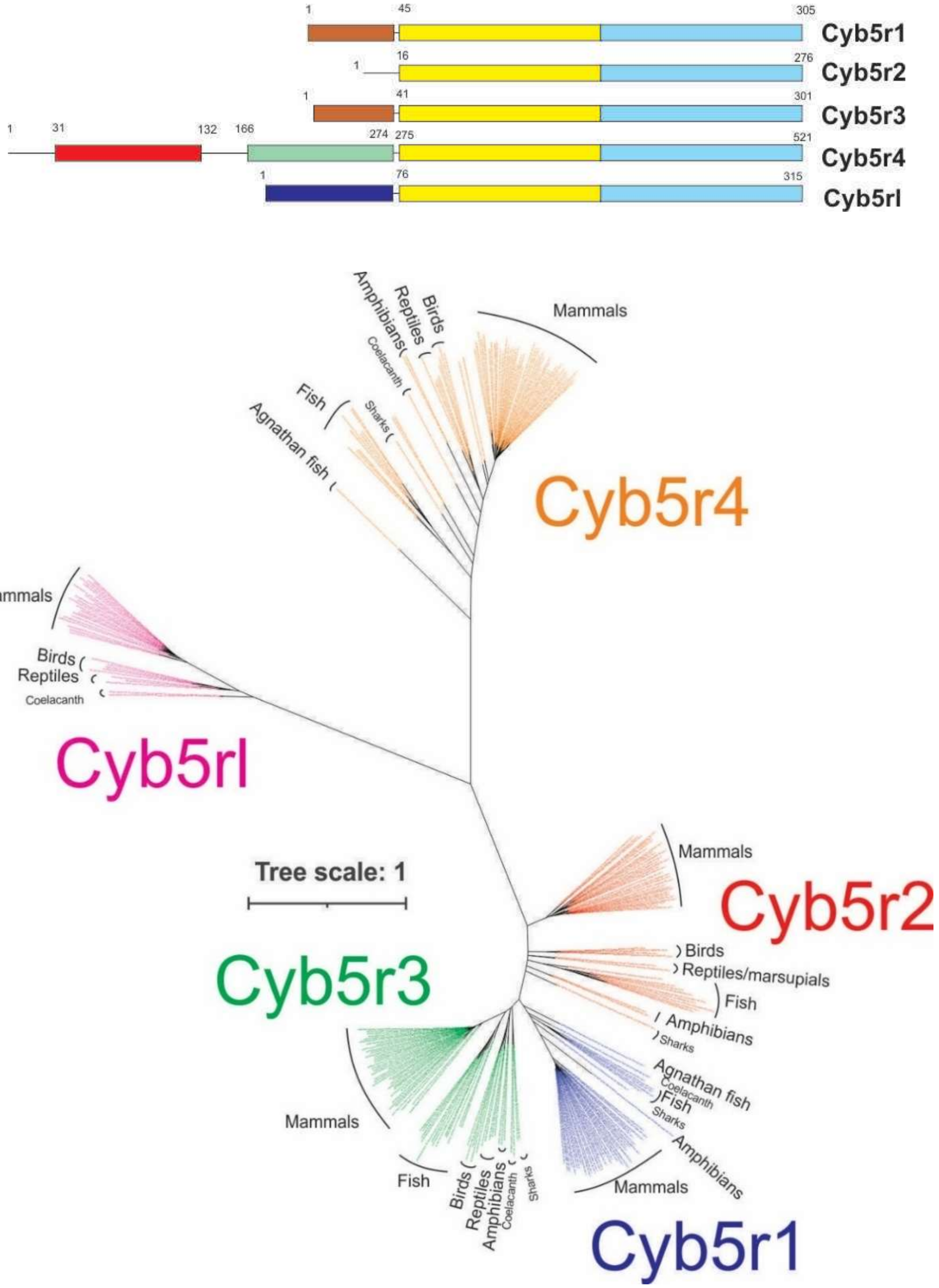

**Supplementary figure 1. Domain arrangement and phylogeny of cytochrome $b_5$ reductases.** Top, domain arrangement in CYB5Rs. CYB5R1/2/3 are highly similar in sequence, with FAD (yellow) and NAD)P)H (light blue) motifs commonly found in the ferredoxin-NADP reductase protein family [30] but differ on short segments in the N-terminus (brown) present in Cyb5R1 and CYb5R3 (but not in CYB5R2) related to mebrane anchoring. CYB5R4 contains a heme binding domain (red) hoologous to cytochrome b5 sequences and linked to the main CYB5R module by a connecting domain (green). CYB5RL contains an additional N terminus sequence (dark blue) different from that found in CYB5R1/2/3 and of unknown function. Numbers indicate amino acid numbers for the human sequence. Bottom, phylogeny of vertebrate CYB5R sequences. The unrooted tree indicates the maximum likelihood distribution of vertebrate CYBR sequences. CYB5R 1/2/3 and 4 are highly conserved in vertebrates; and form two clearly separated clades. CYB5RL is not conserved in vertebrates, being absent in most fish although present in the coelacanth genome.

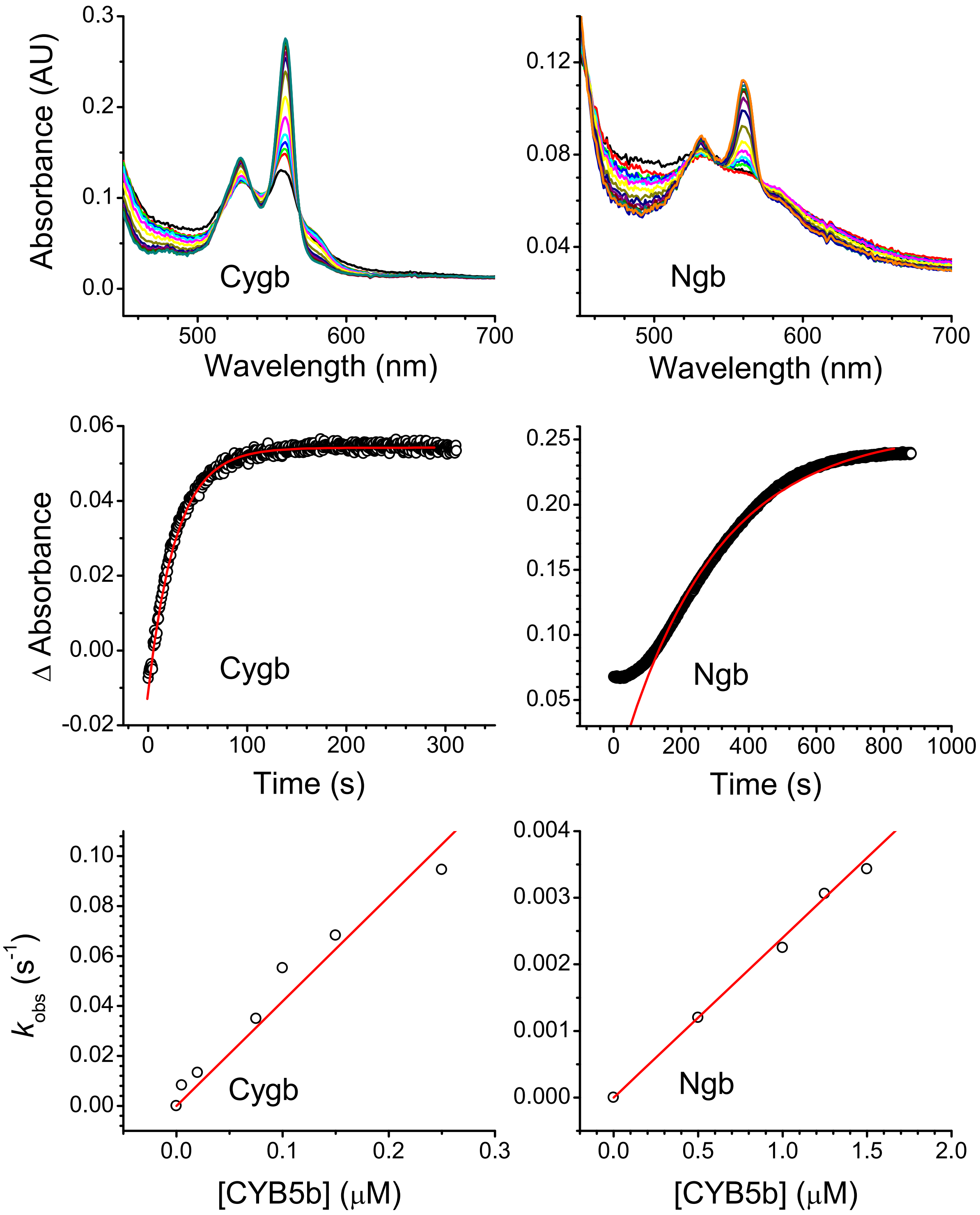

**Supplementary Figure 2. Spectral changes and reaction kinetics for the reaction of CYB5R3/CYB5B with wild type cytoglobin and neuroglobin.** Top left panel shows the reaction of 1 nM CYB5R3, 75nM CYB5B and 5 μM wt Cygb. Top right panel shows the reaction of 1 nM CYB5R3, 1750nM CYB5B and 10 μM wt Ngb. Middel panels show the absorbance changes at selected wavelengths. A lag phase is observed in the first 50 s of reaction until the pool of CYB5B is fully reduced by CYB5R3/NADH. Bottom panels show the observed rates as a function of CYB5b concentration; the slope yields the bimolecular reaction rate constant (CYB5R3 is kept constant at 1nM). Reactions were initiated by the addition of 100 μM NADH. Reactions were carried out in phosphate buffered saline at pH 7.4 and 37°C.

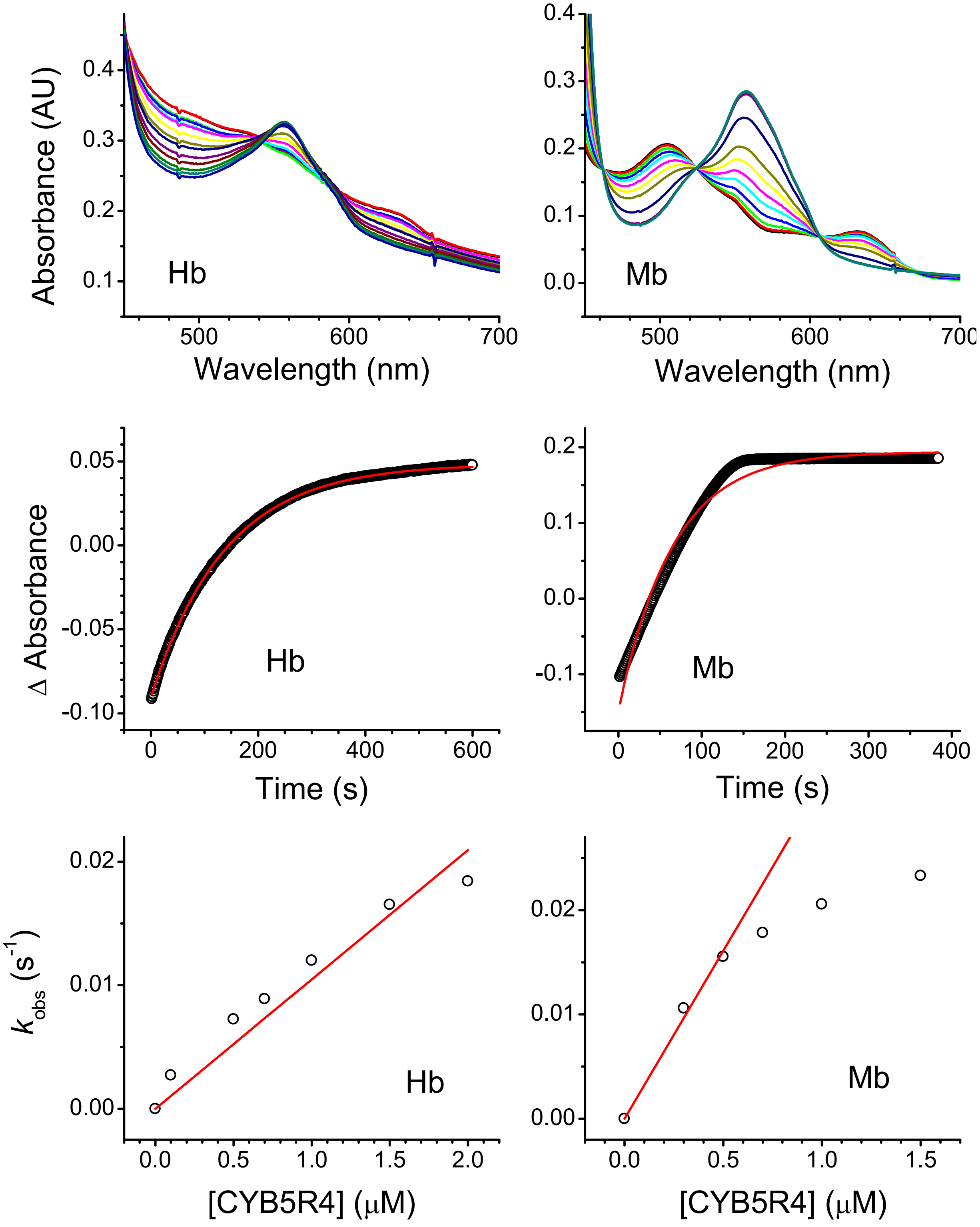

**Supplementary Figure 3. Spectral changes and reaction kinetics for the reaction of CYB5R4 with hemoglobin and myoglobin.** Top left panel shows the reaction of 500 nM CYB5R4, and 20 μM wt Hb. Top right panel shows the reaction of 500 nM CYB5R4 and 20 μM wt Mb. Middel panels show the absorbance changes at selected wavelengths. Bottom panels show the observed rates as a function of CYB5R4 concentration; the slope yields the bimolecular reaction rate constant. Reactions were initiated by the addition of 100 μM NADH. Reactions were carried out in phosphate buffered saline at pH 7.4 and 37°C.

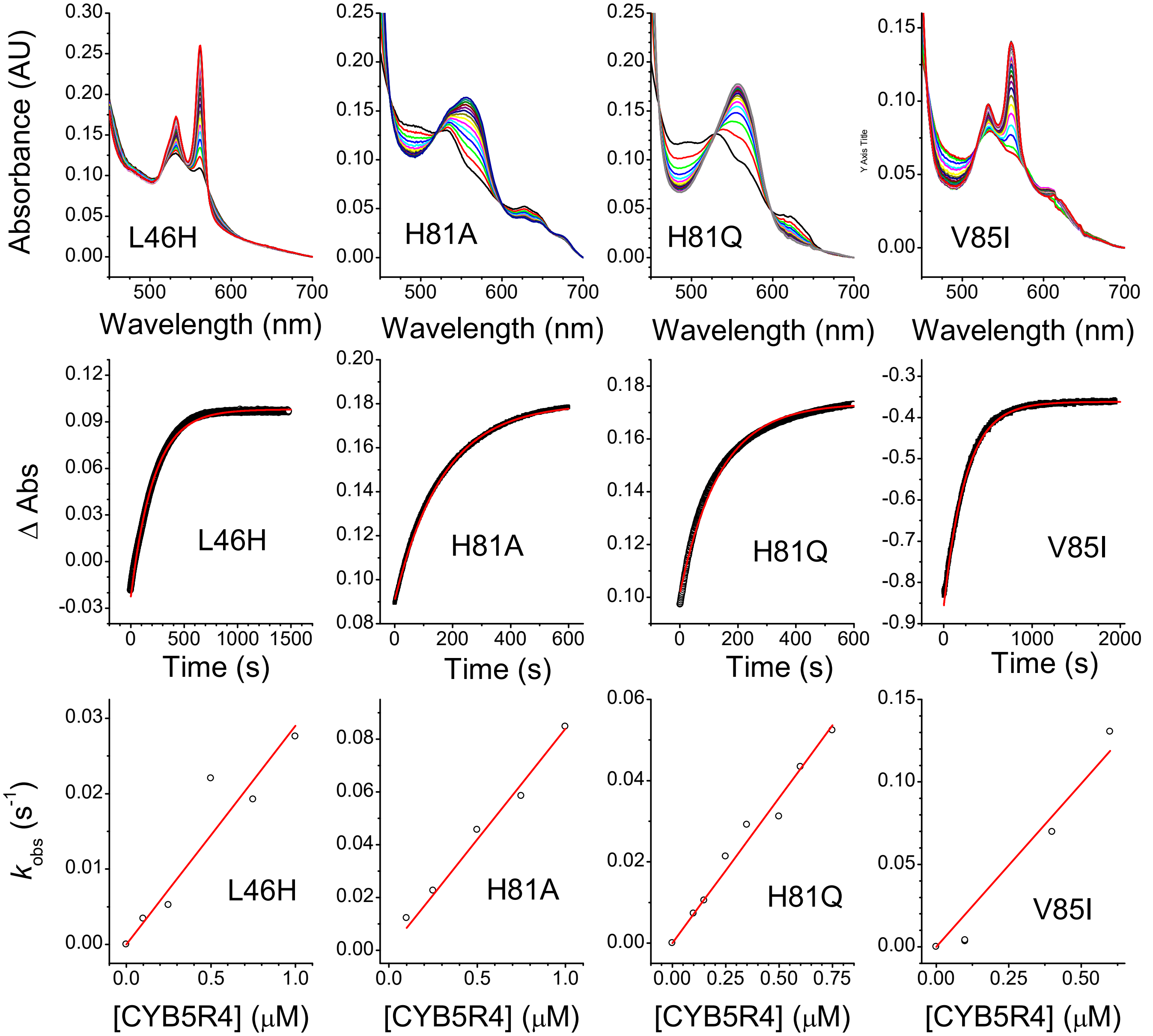

**Supplementary Figure 4.** Spectral changes and reaction kinetics for the reaction of CYB5R4 with cytoglobin mutants L46H, H81A, H81Q, and V85I. Top panels show the spectral changes during the reaction of CYB5R4 with each cytoglobin mutant. Middle panels show the absorbance changes at selected wavelengths. Bottom panels show the observed rates as a function of CYB5R4 concentration; the slope yields the bimolecular reaction rate constant. Reactions were initiated by the addition of 100 μM NADH. Reactions were carried out in phosphate buffered saline at pH 7.4 and 37°C

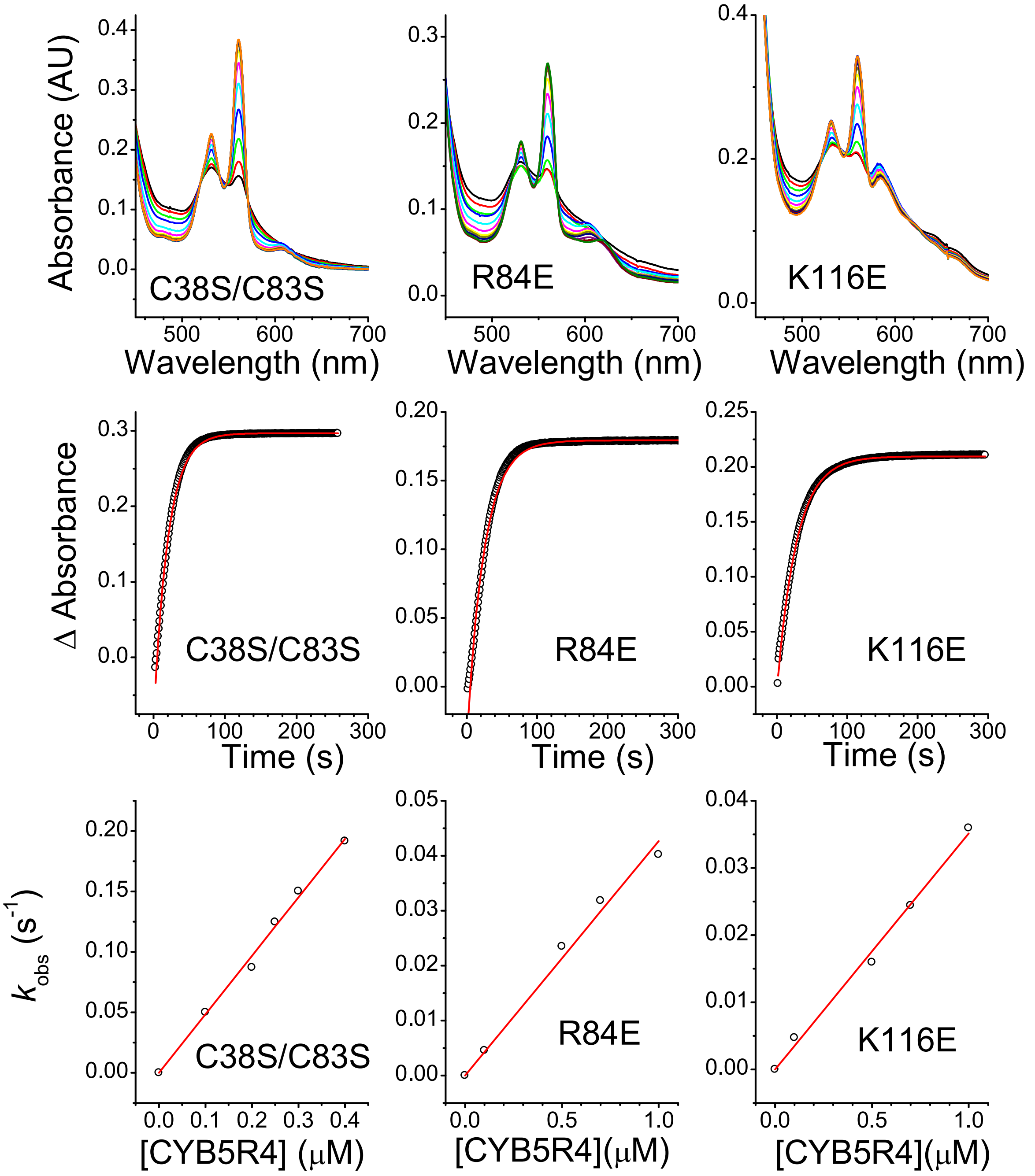

**Supplementary Figure 5. Spectral changes and reaction kinetics for the reaction of CYB5R4 with cytoglobin mutants C38S/C83S, R84E, and K116E**. Top panels show the spectral changes during the reaction of CYB5R4 with each cytoglobin mutant. Middle panels show the absorbance changes at selected wavelengths. Bottom panels show the observed rates as a function of CYB5R4 concentration; the slope yields the bimolecular reaction rate constant. Reactions were initiated by the addition of 100 μM NADH. Reactions were carried out in phosphate buffered saline at pH 7.4 and 37°C

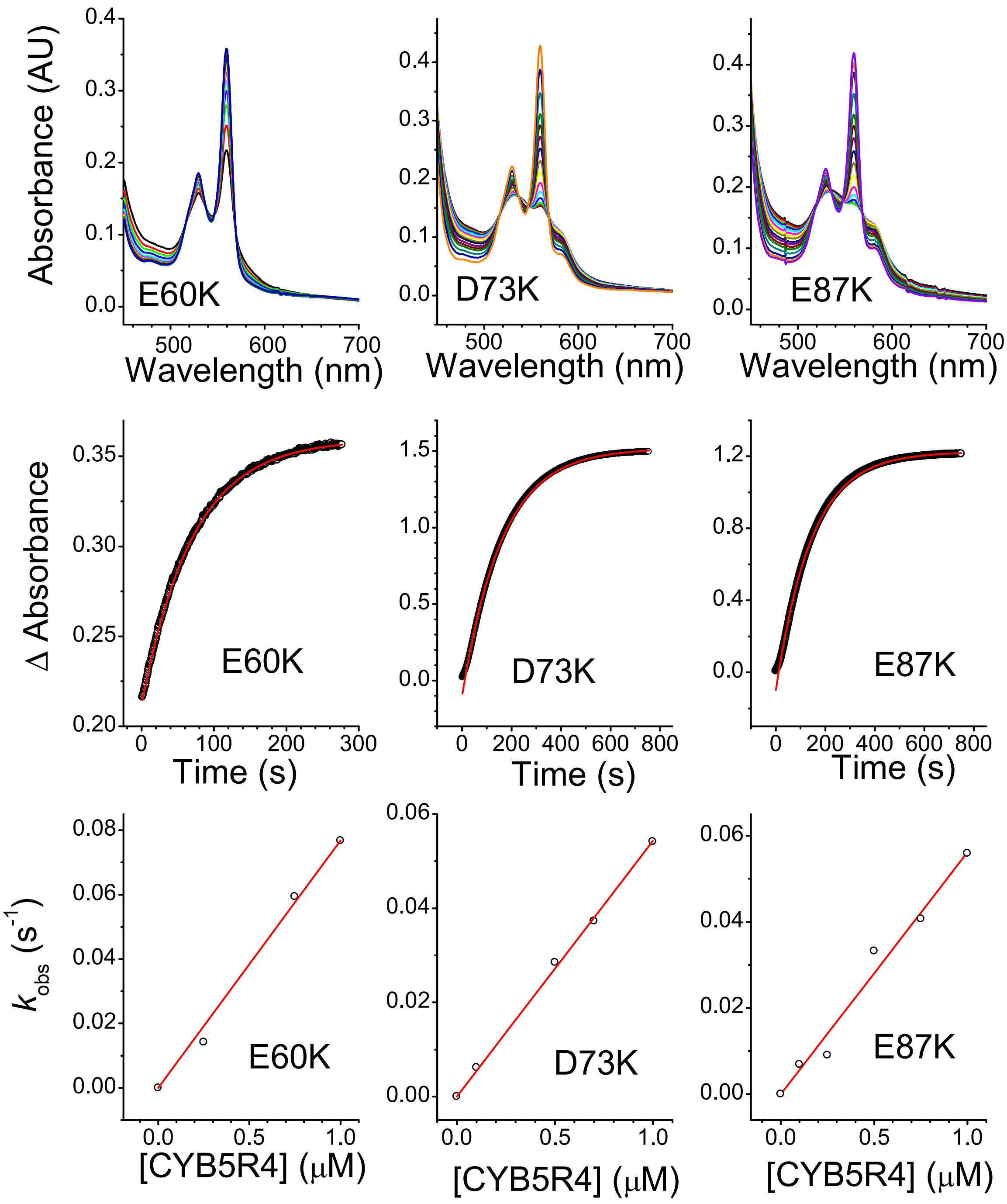

**Supplementary Figure 6. Spectral changes and reaction kinetics for the reaction of CYB5R4 with neuroglobin mutants E60K, D73K, and E87K.** Top panels show the spectral changes during the reaction of CYB5R4 with each neuroglobin mutant. Middle panels show the absorbance changes at selected wavelengths. Bottom panels show the observed rates as a function of CYB5R4 concentration; the slope yields the bimolecular reaction rate constant. Reactions were initiated by the addition of 100 μM NADH. Reactions were carried out in phosphate buffered saline at pH 7.4 and 37°C

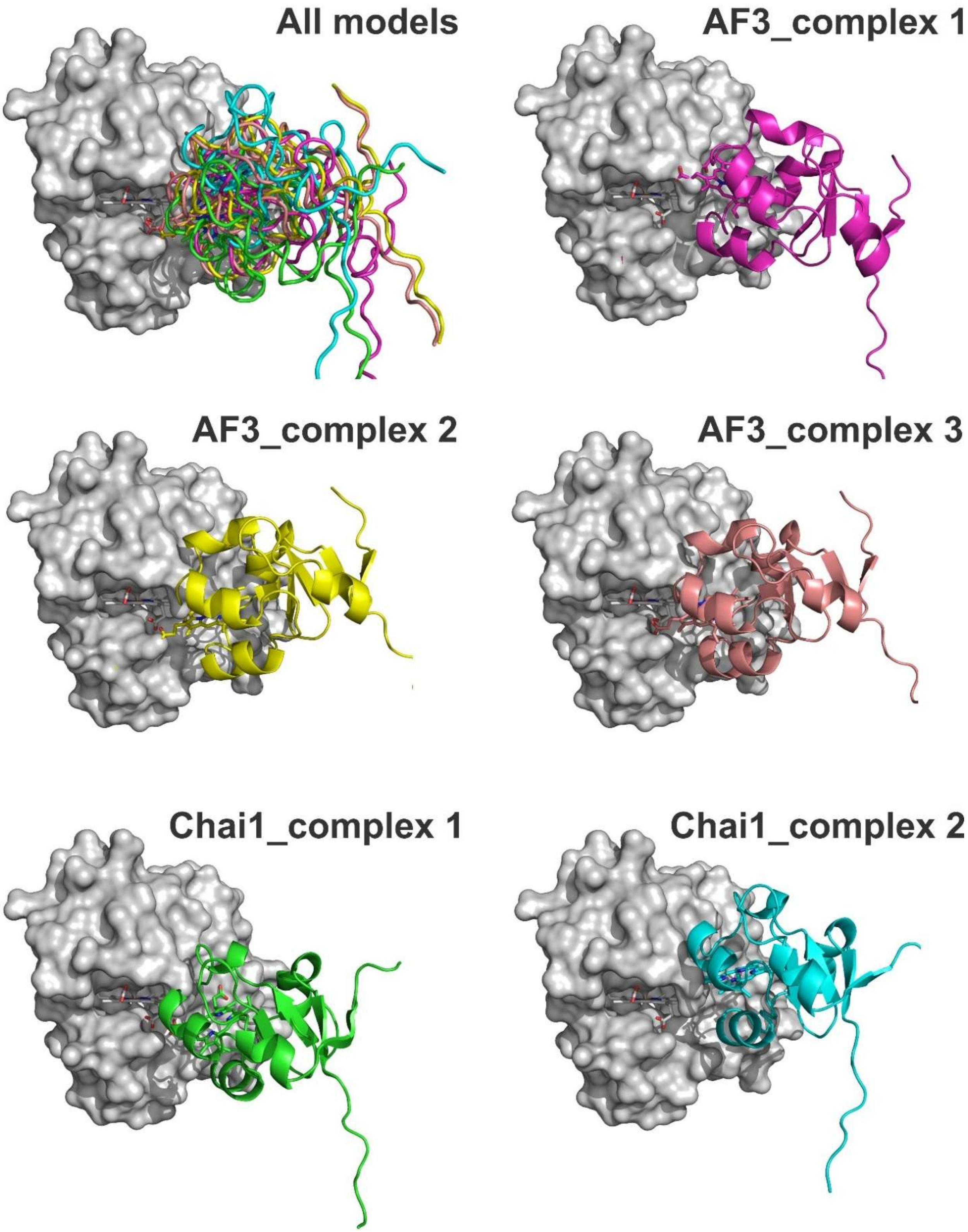

**Supplementary figure 7. Modeled structures for the CYB5R4-Cytoglobin complexes using Alphafold3 and Chai-1.** Only the 5 structures showing optimal heme-heme distances are shown. In all models, Cygb is shown in the same orientation (and same as shown in Figure 5). Cygb molecular surface is shown in light grey, and the Cygb heme moiety is depicted as sticks. The structure of the CYB5R4 CYB5 domain is shown as cartoon except for the combined figure (Top left) where the CYB5R4 secondary structures are shown as ribbons for clarity. Heme groups are shown as sticks. Figure drawn with PyMOL [39].

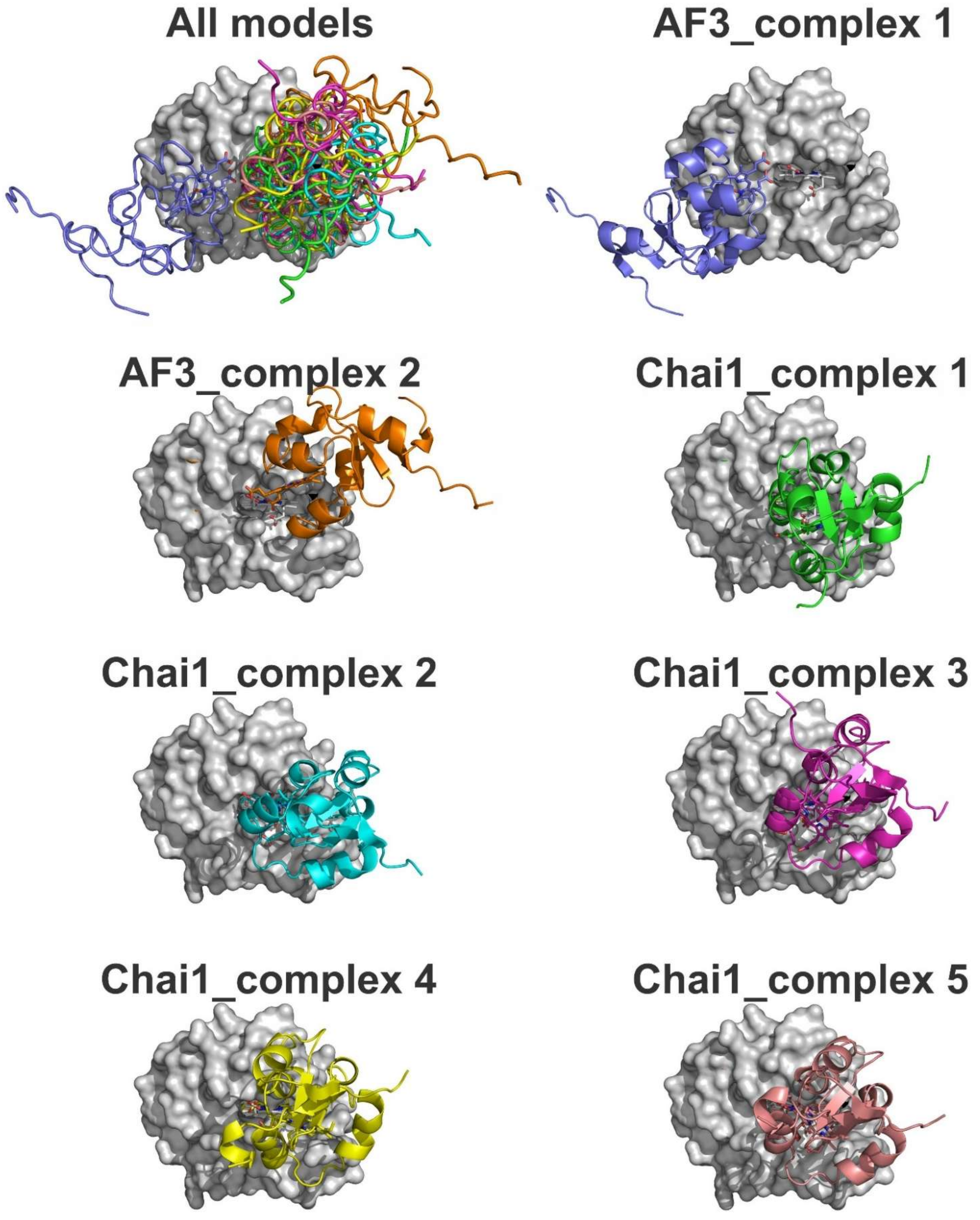

**Supplementary figure 8. Modeled structures for the CYB5R4-Neuroglobin complexes using Alphafold3 and Chai-1.** Only the 7 structures showing optimal heme-heme distances are shown. In all models, Ngb is shown in the same orientation (and same as shown in Figure 5). Ngb molecular surface is shown in light grey, and the Ngb heme moiety is depicted as sticks. The structure of the CYB5R4 CYB5 domain is shown as cartoon except for the combined figure (Top left) where the CYB5R4 secondary structures are shown as ribbons for clarity. Heme groups are shown as sticks. Figure drawn with PyMOL [39].

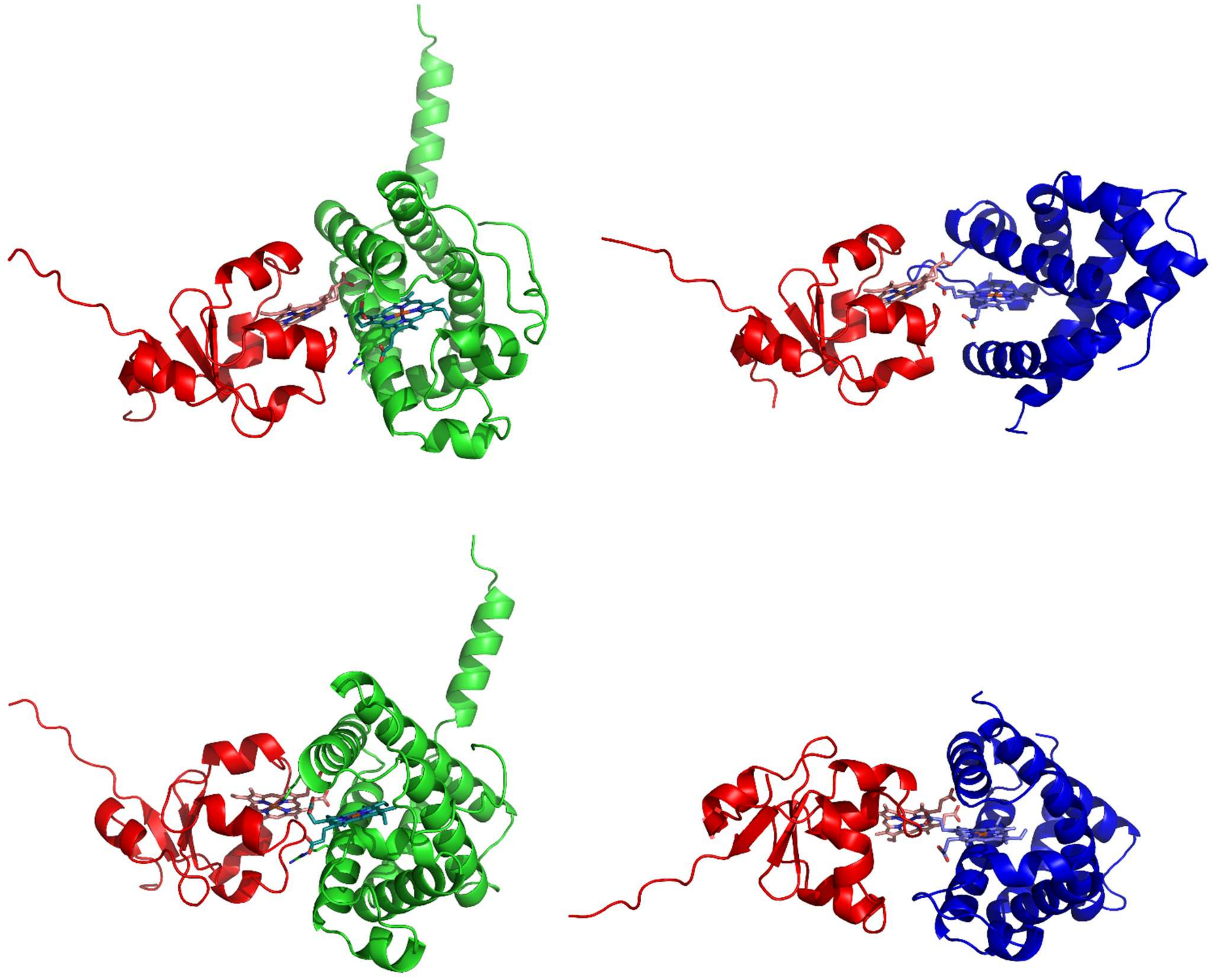

**Supplementary Figure 9. Comparison of selected models for the interaction of the cytochrome *b*$_5$ domain of CYB5R4 with Cytoglobin and Neuroglobin.** The Chai-1 model #1 for the CYB5R4/Cygb and Chai-1 model #1 for the CYB5R4/Ngb interaction are compared. Top panels show the CYB5R4/Cygb model (left) and the CYB5R4/Ngb model (right) showing the CYB5R4 CYB5 domain in the same orientation. Bottom panels show the CYB5R4/Cygb model (left) and the CYB5R4/Ngb model (right) showing the globin in the same orientation. The CYB5R4 cytochrome *b*$_5$ domain (red), Cygb (green) and Ngb (blue) are shown in cartoon representation. Heme moieties are shown as sticks. Figure generated with PyMOL [39].

**Supplementary Table 1. Main properties and interface residues for the selected CYB5R4/Cygb and CYB5R4/Ngb complexes**

| **Cygb-Cb5r4 complexes** | | | | | | | | | |
|---|---|---|---|---|---|---|---|---|---|
| **AF complex 1** | | **AF complex 2** | | **AF complex 3** | | **Chai1 complex 1** | | **Chai1 complex 2** | |
| Binding surface (Avge) = 414.7 Å$^2$ | | Binding surface (Avge) = 582.2 Å$^2$ | | Binding surface (Avge) = 563.5 Å$^2$ | | Binding surface (Avge) = 409.7 Å$^2$ | | Binding surface (Avge) = 329.5 Å$^2$ | |
| Heme-heme distance = 5.80 Å | | Heme-heme distance = 3.01 Å | | Heme-heme distance = 2.23 Å | | Heme-heme distance = 3.09 Å | | Heme-heme distance = 3.09 Å | |
| **Surface residues** | | **Surface residues** | | **Surface residues** | | **Surface residues** | | **Surface residues** | |
| **Cygb** | **Cb5r4** | **Cygb** | **Cb5r4** | **Cygb** | **Cb5r4** | **Cygb** | **Cb5r4** | **Cygb** | **Cb5r4** |
| Tyr35 | Arg76 | Tyr35 | Arg76 | Tyr35 | | Tyr35 | | Tyr35 | |
| | | | Tyr85 | | Tyr85 | | | Cys38 | |
| | | Glu39 | Met86 | Glu39 | Met86 | | Met86 | Glu39 | |
| | | | Glu87 | | Glu87 | | Glu87 | Pro76 | Glu87 |
| | | | Tyr88 | Arg79 | Tyr88 | | Tyr88 | | |
| Lys80 | | Lys80 | His89 | Lys80 | His89 | Lys80 | His89 | Lys80 | His89 |
| Cys83 | Pro90 | Cys83 | Pro90 | Cys83 | Pro90 | Cys83 | Pro90 | Cys83 | Pro90 |
| Arg84 | Gly91 | Arg84 | Gly91 | Arg84 | Gly91 | Arg84 | Gly91 | Arg84 | Gly91 |
| | | | Gly92 | | Gly92 | | Gly92 | Met86 | Gly92 |
| Gly87 | | Gly87 | Glu93 | Gly87 | Glu93 | Gly87 | Glu93 | Gly87 | |
| Ala88 | | Ala88 | | Ala88 | | Ala88 | Asp94 | | Asp94 |
| Asn90 | | Asn90 | | Asn90 | | | Glu95 | | Glu95 |
| Thr91 | | Thr91 | | Thr91 | | Thr91 | | Thr91 | Arg98 |
| Glu94 | Gln110 | Glu94 | | Glu94 | Gln110 | | Gln110 | | |
| Asn95 | Val111 | Asn95 | Val111 | Asn95 | Val111 | | Val111 | Asn95 | Val111 |
| | | | | | | Asp98 | His112 | | |
| | Arg113 | | Arg113 | | Arg113 | Asp100 | | | Arg113 |
| Lys101 | Trp114 | Lys101 | Trp114 | Lys101 | Trp114 | Lys101 | Trp114 | Lys101 | Trp114 |
| Ser104 | Val115 | Ser104 | Val115 | Ser104 | Val115 | Ser104 | Val115 | Ser104 | |
| Val105 | Asn116 | Val105 | Asn116 | Val105 | Asn116 | Val105 | | Val105 | |
| Leu108 | Tyr117 | Leu108 | | Leu108 | | Leu108 | | Leu108 | |
| Val109 | Glu118 | Val109 | | Val109 | | | | | |
| Ala112 | Ser119 | | Ser119 | | Ser119 | | | | |
| Lys116 | Met120 | | Met120 | Lys116 | Met120 | Lys116 | | | |

**Supplementary Table 1 (continued)**

| **Ngb-Cb5r4 complexes** | | | | | | | | | | | | | |
|---|---|---|---|---|---|---|---|---|---|---|---|---|---|
| **AF complex 1** | | **AF complex 2** | | **Chai1 complex 1** | | **Chai1 complex 2** | | **Chai1 complex 3** | | **Chai1 complex 4** | | **Chai1 complex 5** | |
| Binding surface (Avge) = 323.2 Å$^2$ | | Binding surface (Avge) = 535.2 Å$^2$ | | Binding surface (Avge) = 271.1 Å$^2$ | | Binding surface (Avge) = 397.4 Å$^2$ | | Binding surface (Avge) = 376.2 Å$^2$ | | Binding surface (Avge) = 298.0 Å$^2$ | | Binding surface (Avge) = 345.4 Å$^2$ | |
| Heme-heme distance = 4.10 Å | | Heme-heme distance = 2.55 Å | | Heme-heme distance = 3.26 Å | | Heme-heme distance = 2.97 Å | | Heme-heme distance =3.36 Å | | Heme-heme distance = 3.27 Å | | Heme-heme distance = 2.00 Å | |
| **Surface residues** | | **Surface residues** | | **Surface residues** | | **Surface residues** | | **Surface residues** | | **Surface residues** | | **Surface residues** | |
| **Ngb** | **Cb5r4** | **Ngb** | **Cb5r4** | **Ngb** | **Cb5r4** | **Ngb** | **Cb5r4** | **Ngb** | **Cb5r4** | **Ngb** | **Cb5r4** | **Ngb** | **Cb5r4** |
| Ser17 | | Pro40 | | | | | | | Lys68 | | | | |
| Pro20 | Arg76 | Leu41 | Arg76 | | | | | | | Leu41 | | Leu41 | |
| | | Gln43 | | | | Gln43 | Pro84 | Gln43 | | Gln43 | | Gln43 | |
| | | Tyr44 | Tyr85 | | | Tyr44 | Tyr85 | Tyr44 | Tyr85 | Tyr44 | Tyr85 | Tyr44 | Tyr85 |
| | | Asn45 | | Asn45 | Met86 | Asn45 | | Asn45 | | Asn45 | | Asn45 | |
| | | Cys46 | Glu87 | Cys46 | Glu87 | Cys46 | Glu87 | Cys46 | Glu87 | Cys46 | | Cys46 | |
| | | | Tyr88 | | Tyr88 | Arg47 | Tyr88 | | Tyr88 | | | | Tyr88 |
| | | | His89 | | His89 | Gln48 | His89 | Gln48 | His89 | | | | His89 |
| | | | Pro90 | Glu60 | Pro90 | | Pro90 | | Pro90 | | Pro90 | | Pro90 |
| Arg66 | | | | | Gly91 | | Gly91 | | Gly91 | | Gly91 | | Gly91 |
| | | Lys67 | | Lys67 | Gly92 | | Gly92 | | Gly92 | | Gly92 | Lys67 | Gly92 |
| Leu70 | | | | | Glu93 | | Glu93 | | Glu93 | | Glu93 | | Glu93 |
| Val71 | | | | | | | Asp94 | | Asp94 | | Asp94 | | Asp94 |
| Asp73 | | | | | | | Glu95 | | Glu95 | | Glu95 | | Glu95 |
| Ala74 | | | | | Val111 | | | | | | Val111 | | Val111 |
| Thr77 | | | | | His112 | | | | | | His112 | | |
| Asn78 | Arg113 | | Arg113 | | | | | | | | | | |
| Glu87 | Trp114 | | Trp114 | | Trp114 | | | | | | Trp114 | | |
| Tyr88 | Val115 | | Val115 | | Val115 | | | | Val115 | | Val115 | | Val115 |
| | Asn116 | Arg94 | Asn116 | | Asn116 | | | Arg94 | | Arg94 | | Arg94 | |
| | Ser119 | Lys95 | Ser119 | Lys95 | | Lys95 | | Lys95 | | Lys95 | | Lys95 | |
| | Met120 | Arg97 | Met120 | | Met120 | | Met120 | | Met120 | | Met120 | | Met120 |
| | | Ala98 | | Ala98 | | Ala98 | Leu121 | Ala98 | | Ala98 | | Ala98 | |
| | | Val99 | | Val99 | | Val99 | Cys124 | Val99 | | Val99 | | Val99 | |
| | | Gly100 | | | | | | | | | | | |
| | | Gly150 | | | | | | | | | | | |
